# Wind Gates Olfaction Driven Search States in Free Flight

**DOI:** 10.1101/2023.11.30.569086

**Authors:** S. David Stupski, Floris van Breugel

## Abstract

For organisms tracking a chemical cue to its source, the motion of their surrounding fluid provides crucial information for success. Swimming and flying animals engaged in olfaction driven search often start by turning into the direction of an oncoming wind or water current. However, it is unclear how organisms adjust their strategies when directional cues are absent or unreliable, as is often the case in nature. Here, we use the genetic toolkit of *Drosophila melanogaster* to develop an optogenetic paradigm to deliver temporally precise “virtual” olfactory experiences for free-flying animals in either laminar wind or still air. We first confirm that in laminar wind flies turn upwind. Furthermore, we show that they achieve this using a rapid (∼100 ms) turn, implying that flies estimate the ambient wind direction prior to “surging” upwind. In still air, flies adopt remarkably stereotyped “sink and circle” search state characterized by ∼60°turns at 3-4 Hz, biased in a consistent direction. Together, our results show that *Drosophila melanogaster* assess the presence and direction of ambient wind prior to deploying a distinct search strategy. In both laminar wind and still air, immediately after odor onset, flies decelerate and often perform a rapid turn. Both maneuvers are consistent with predictions from recent control theoretic analyses for how insects may estimate properties of wind while in flight. We suggest that flies may use their deceleration and “anemometric” turn as active sensing maneuvers to rapidly gauge properties of their wind environment before initiating a proximal or upwind search routine.

## INTRODUCTION

The ability to track a chemical cue to its source is critical for the fitness of motile organisms with diverse life histories. Bacteria with no brain navigate to the source of nutrients^1^, benthic isopods track the smell of carrion on the dark ocean floor^2^, and moths locate nectar-laden flowers in complex chemical landscapes^3^. To succeed in tracking a plume to its source animals must contend with cues that are intermittently dispersed in space by natural advection processes^4^. Over a century of research^5,6^ has provided a clear picture of how animals track plumes in consistent laminar flow. A nearly ubiquitous strategy is to adopt a stereotyped “cast and surge” behavior; surging into the direction of flow upon encountering a chemical plume and performing cross flow, casting counter turns after losing track of the plume^7–11^.

Cast and surge has served as the scaffolding for ethological, robotic, and algorithmic studies of plume tracking since the turn of the 20th century^5,12^. However it is unclear whether cast and surge represents a one size fits all sensorimotor motif for successful plume tracking, or if there are alternative search states which optimize successful plume tracking depending on the flow conditions. For flying insects, both elements of cast and surge rely on a directional cue from the wind, but in nature the wind direction can shift on timescales relevant for plume tracking, or be very still^13–15^. To address this knowledge gap we focus on the extreme scenario with still air, and ask a simple but deceptively complex question: what sensorimotor transformations underlay search when there is no directional information from the wind? To answer this question we developed an optogenetic paradigm that allows spatiotemporally precise delivery of a fictive odor plume independent of the wind environment for free-flying *Drosophila melanogaster*. In still air, flies adopt a previously undescribed, highly stereotyped search behavior characterized by a sequence of rapid turns in a consistent direction while also lowering their altitude. This “sink and circle” style of plume tracking is the still air complement to the well described cast and surge routine that flies perform in the presence of laminar wind. Furthermore, we reveal a previously overlooked aspect of the cast and surge strategy in *Drosophila*. Flies surge into the wind with kinematics closely matching those of body saccades, rapid (∼100 ms) feed-forward motor programs^16,17^.

Our discovery of sink and circle, and the saccadic upwind turn, indicates that flies can both estimate the presence and direction of ambient wind while flying. However, without access to stationary sensors, or magnitude calibrated sensory systems, a flying insect cannot directly measure information about the wind environment^18,19^. Instead, they perceive apparent airflow, the vector sum of ambient wind and the relative airflow due to self-motion. Historically, wind orientation in insect flight has been attributed to visual anemotaxis^20,21^: a wind orienting strategy that primarily uses angular measurements of ventral optic flow and the apparent airflow direction to guide an insect into the direction of ambient wind. The only stable solution to a control strategy that minimizes the difference between these two angles is to orient upwind^22^. Visual anemotaxis, however, would not allow a fly to distinguish between flying upwind or flying in still air, nor would it explain how a fly can perform a directed saccade into the direction of ambient wind, or maintain a crosswind casting course. Our results indicate that flying *Drosophila* do not rely on visual anemotaxis to turn into the wind. Instead, our data offer support to an alternative hypothesis^22^: that flies use an active sensing maneuver consisting of decelerating and a rapid change in course direction at the onset of an odor event to gauge properties of the wind.

## RESULTS

### Activating *Orco+* neurons in flight generates stereotypical cast and surge behaviors in laminar wind

To independently control the wind and olfactory experience of flying flies we developed a custom-built wind tunnel capable of delivering 40 cm/s laminar wind or still air (Figure 1A). We expressed the red-light sensitive ion channel, CsChrimson^23^, in the expression pattern of Orco, a co-receptor necessary for proper function of a broad group of olfactory receptors^24^. The tunnel was equipped with machine vision cameras to track flies in 3-D^25^ at 100 Hz against uniform infrared illumination, while flies were provided with a combination of blue and white illumination (425 lux total) so they could navigate the space. When a fly entered a pre-defined “trigger zone” (Figure 1B) it was assigned either a “flash” condition which turned on two symmetric rows of red LEDs for 675 ms (42 *µ* W *mm*^−2^, Figure S1), or a “sham” corresponding to no light flash (Figure 1B).

**Figure 1.**
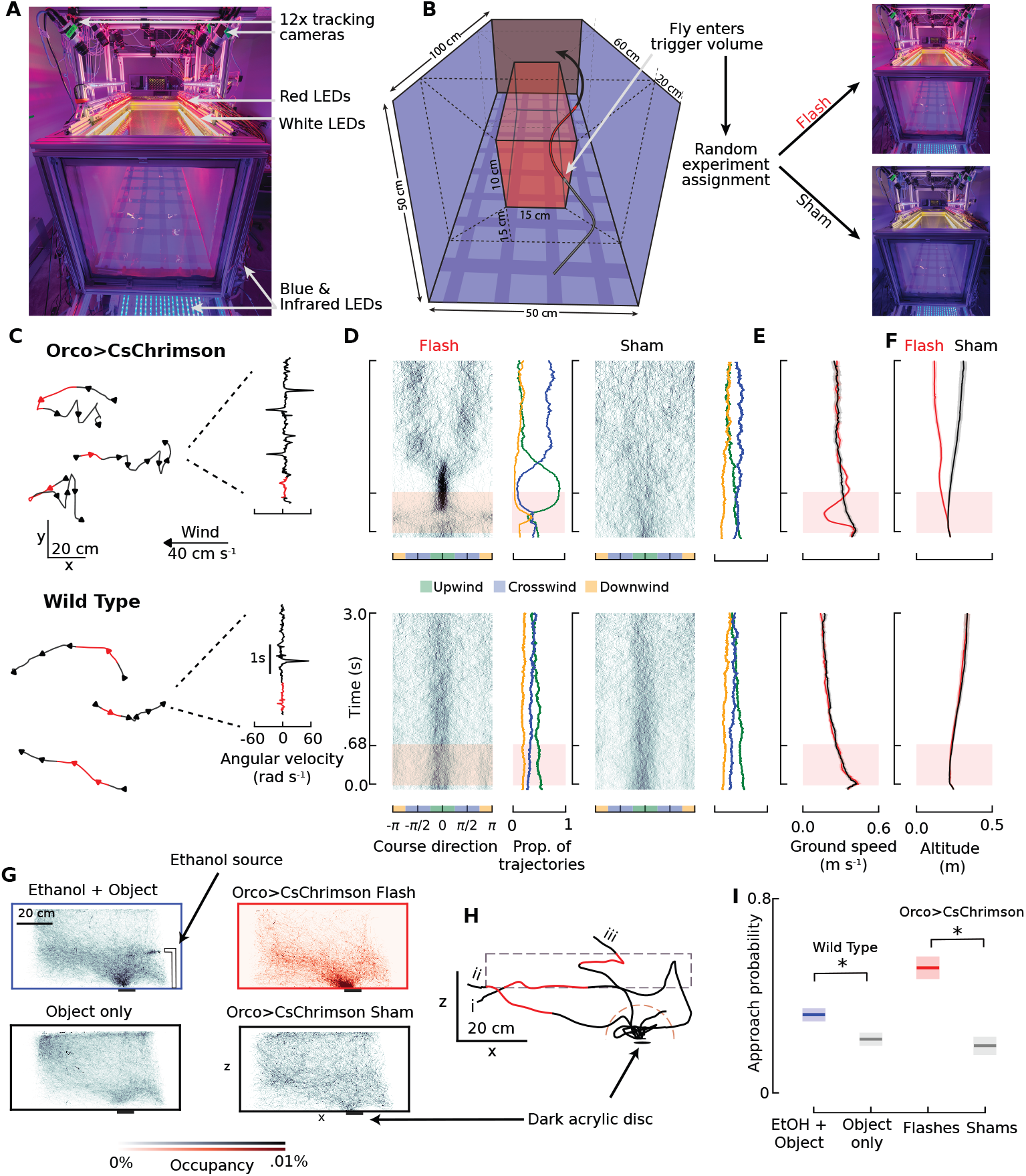
Eliciting stereotyped plume-tracking responses using optogenetics. (A) Wind tunnel system for free-flight optogenetics. (B) Experimental design: a trajectory enters the trigger zone and is randomly assigned a “flash” or “sham” event. (C) Three sample trajectories in the x, y plane of Orco>CsChrimson flies and wild-type flies and their response to a 675 ms flash of red light. Red regions indicate the portions of trajectories during which the 675 ms flash was activated. (D) Course direction vs time relative to the trajectory entering the trigger zone for Orco>CsChrimson flies (top) and wild-type flies (bottom) that received either a flash stimulus or a sham event (Orco>CsChrimson N=60 flies: flash, n=232 trajectories; Orco>CsChrimson, sham n=222 trajectories; wild-type N=75 flies: flash n=398 trajectories; wild-type, sham n= 429 trajectories). A course direction of 0 indicates upwind and *π*/-*π* is downwind. Next to each course direction heat map is the fraction of trajectories oriented upwind (green), crosswind (blue) or downwind (orange). (E) Ground speed response of Orco>CsChrimson and wild-type flies in response to a red light stimulus (red trace) compared to shams (black trace). (F) Same as (E) except for altitude responses from optogenetic stimulus. (G) Heat maps in the x, z plane of trajectory occupancies within the wind tunnel volume with a dark feature on the tunnel floor. (H) Three example trajectories of Orco> CsChrimson flies entering the trigger zone (dashed rectangle), receiving a stimulus, and entering the airspace above the object (dashed semicircle). (I) Object approach probability based on experimental conditions: Wild-type flies with an ethanol plume. N=75 flies: n=1169 trajectories; wild-type flies in clean air N=75 flies: n=856 trajectories; Orco>CsChrimson N=60 flies: trajectories randomly assigned a flash event, n=445; sham events, n=435. Shading in panels throughout the manuscript, indicates 95% confidence intervals. Lines on graphs represent means and error shading on graphical representations of data represent 95% confidence intervals. See also Figures S1, S2 and Video S1

After optogenetic activation of *Orco+* neurons individual flies oriented upwind on average 382 ms (± 103 ms STD) after the stimulus onset (compared to previous^8^ sensorimotor delay estimates of 200 ms) (Figure 1C-D, Figure S2, Video S1). We defined the time of anemotaxis as the first point at which a 50 ms rolling average of the course direction transitioned from greater than 20° off of upwind to less than 20° off of upwind. The time of 382 ms reflects the sum of computational delay of determining the wind direction and the biomechanical delay of anemotaxis itself. In 77% of trajectories we were able to identify a specific time of anemotaxis using this approach. The remaining 23% of flies either maintained an upwind course throughout this entire period, or did not turn into the wind (mean course error from upwind of 15.7^°^ ± 24.5°, Figure 1D). In contrast, we did not find evidence of anemotaxis in sham controls for Orco>CsChrimson flies, flash and sham conditions for wild-type flies (Figure 1C-D), or for a genetic control (UAS-CsChrimson/+, Figure S2). There was however, a barely detectable and time delayed anemotaxis response in Orco>CsChrimson flies raised without supplemental ATR (Figure S2). Approximately 540 ms after the optogenetic stimulus ended, the course direction of Orco>CsChrimson flies was largely perpendicular to the wind direction, characteristic of the casting behaviors observed for insects interacting with a real odor plume^8,26^. These response times are comparable to the 450 ms delay seen in casting initiation after losing contact with a real ethanol plume^8^. We did not see any evidence of casting as a result of the red light flash in any genetic or positional controls (Figure 1D, Figure S2).

Orco>CsChrimson flies displayed a significant drop in ground speed (ground speed throughout refers to speed in the x,y plane) in response to optogenetic stimulation compared to sham trials (Figure 1E, Mann-Whitney U test, U=10436, p=2.9×10^−28^). We found no evidence of this effect in wild-type, or genetic controls, suggesting that it was not a visually mediated startle response. Although this deceleration behavior was not originally reported, our re-analysis of flies tracking a well-defined ethanol plume revealed a similar, albeit more subtle, deceleration effect (Figure S2). Decelerating in response to encountering an odor has also been reported for flies elsewhere^27^.

Orco>CsChrimson flies also reduced their altitude in response to optogenetic activation of *Orco*+ neurons 240 ms after a triggering event (Figure 1F, Mann-Whitney U test,U=14804, p=2.4 ×10^−15^) and continued to maintain a lower altitude throughout the entire time course. The altitude difference levelled out at around 19 cm within 2 seconds of flies entering the trigger zone. Initially we hypothesized that the altitude lowering may have been due to attraction to the visually salient grid. However, when we performed the same experiment without a visual checkerboard on the wind tunnel floor, flies still reduced their altitude in response to optogenetic activation of *Orco*+ neurons (Figure S3). We did not find any noteworthy altitude modulation effects in wild-type flies, or any genetic controls in response to a flash of red light.

### Optogenetic activation of *Orco*+ neurons elicits search for visually contrastive features

Activating *Orco*+ neurons in flight also evoked visually guided search in *Drosophila* similar to published free flight experiments performed with ethanol, vinegar, and CO_2_ plumes^8,28^. We first validated visually evoked search behaviors in our wind tunnel, using an ethanol plume and a 3.8 cm diameter, dark, infrared transmissive, acrylic disc, 35 cm downwind of the odor source. We quantified the probability that a trajectory from wild-type flies approached the object both in the presence and absence of the ethanol plume (Figure 1G). We considered a trajectory to have approached the object if it entered a 10 cm radius around the acrylic disc (Figure 1H).

Even though there was no red light triggering in our wild-type experiments we only included trajectories from flies which entered the trigger zone volume of the tunnel. Consistent with prior work^8^, we found that flies were approximately 50% more likely to enter the airspace around the dark visual feature in the presence of an ethanol plume (Figure 1I, two-proportion Z-test, Z=5.0, p=2.8×10^−7^). Next we used the same optogenetic activation paradigm described in the previous section, only this time with the same dark visual feature used in our ethanol plume experiments. We focused our analysis on the first 5 seconds after a trajectory initiated a triggering event. Orco>CsChrimson trajectories that received a flash of red light approached the object 51.5% of the time, more than twice that when compared to the trajectories assigned a sham event (Figure 1I, two-proportion Z-Test, Z=9.9, p = 3.6×10^−23^). Orco>CsChrimson trajectories from sham events and wild-type flies flying in clean air had similar levels of attraction to the object (Figure 1I, two-proportion Z-test, Z=1.0, p=.14), suggesting that only a very recent experience with an odor elicits visual approach.

### Flies change search states in still air

After establishing that our optogenetic approach elicits the full suite of established olfactory navigation behaviors for flies flying in laminar wind, we set out to answer a question that would not be feasible using a real odor. How do flies modulate their search behavior depending on the presence or absence of wind? Whereas the first few hundred ms after the onset of the olfactory stimulus was similar regardless of whether or not flies were flying in wind or still air, after the stimulus ended trajectories displayed a strikingly different, and highly stereotyped search behavior. Rather than surging and casting, flies in still air performed a sequence of rhythmic uni-directional turns resulting in a tight circling behavior centered around the point where the optogenetic stimulus ended (Figure 2A-B, Videos S2-14). Flies did not orient into any particular part of the wind tunnel (Figure 2C). However, similar to activation of *Orco*+ neurons in wind, flies also sharply decelerated on the stimulus onset (Figure 2D). We did not robustly observe this behavior, or any other artifacts, in either the sham trajectories for Orco>CsChrimson flies or any wild-type controls (Figure S4). The circling behavior consisted of two primary motifs: sharp, regular saccades in a consistent direction, while also reducing altitude.

**Figure 2.**
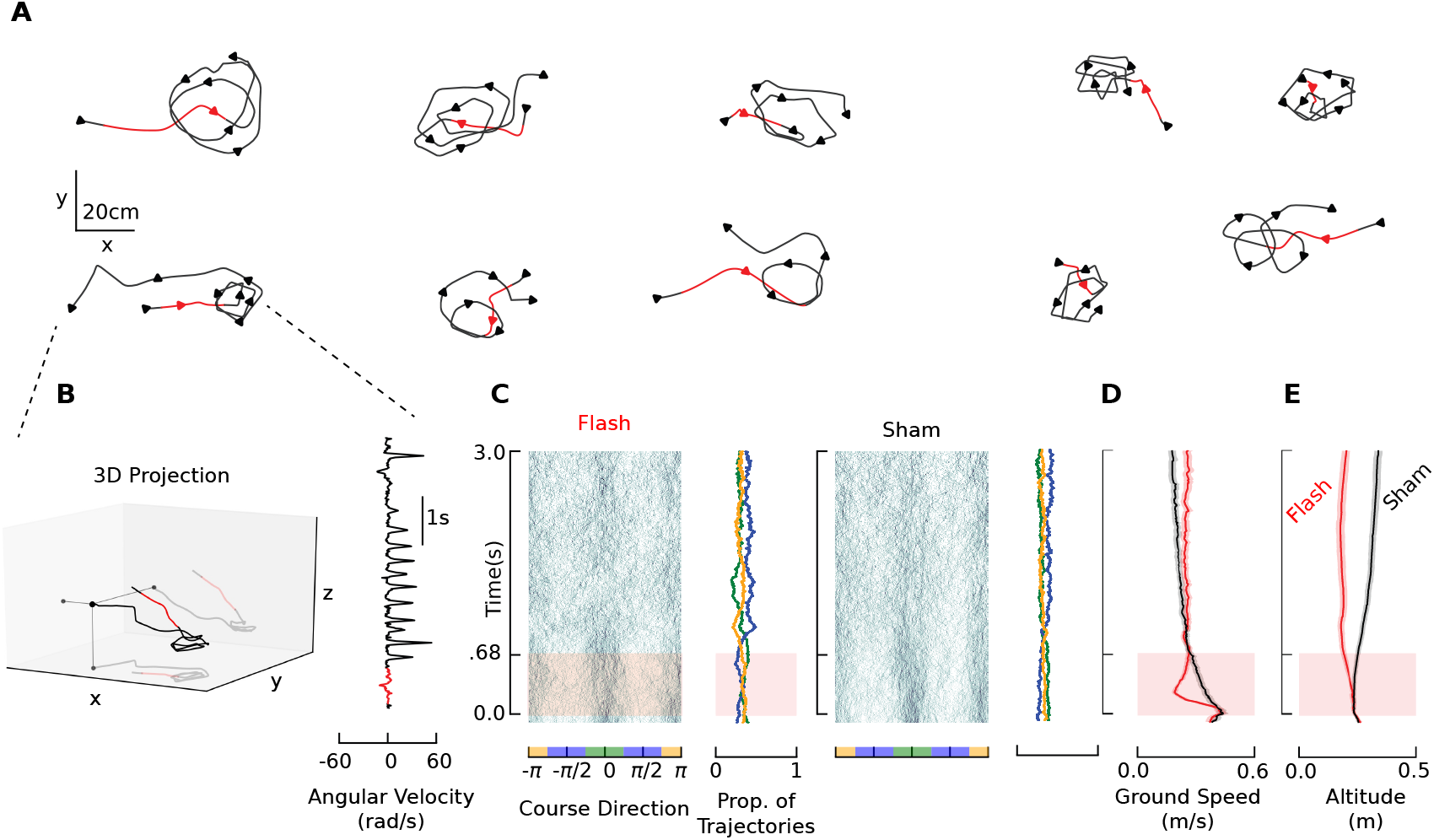
Optogenetic activation of *Orco+* neurons in still air elicits circling rather than casting. (A) Ten sample trajectories of the search behaviors of Orco>CsChrimson flies following optogenetic stimulation of *Orco*+ neurons in still air. (B) Sample 3D projection and angular velocity profile of a trajectory. (C) Course direction in the same reference frame as Figure 1D. N=60 flies, n=263 flash event trajectories, and n=291 sham event trajectories. (D) Ground speed changes in response to activating of *Orco*+ neurons in the absence of wind. (E) Altitude responses of Orco>CsChrimson flies in response to an optogenetic stimulus in the absence of wind. Lines on graphs represent means and error shading on graphical representations of data represent 95% confidence intervals. See also Figures S3-4 and Videos S2-14.

### Circling is Characterized by Modulating the Amplitude, Frequency, and Directional Bias of Saccades

Flying fruit flies typically change course via rapid turns referred to as saccades in the literature^16,29–31^. To quantify the stark differences in post-stimulus saccade dynamics in laminar wind vs. still air, we use a slightly modified Geometric Saccade Detection algorithm originally found in^30^ (Figure 3A, Figure S5) and characterized properties of saccades identified from 0.9-3 s after the stimulus. This allowed us to compare saccades specifically from circling and casting behaviors.

**Figure 3.**
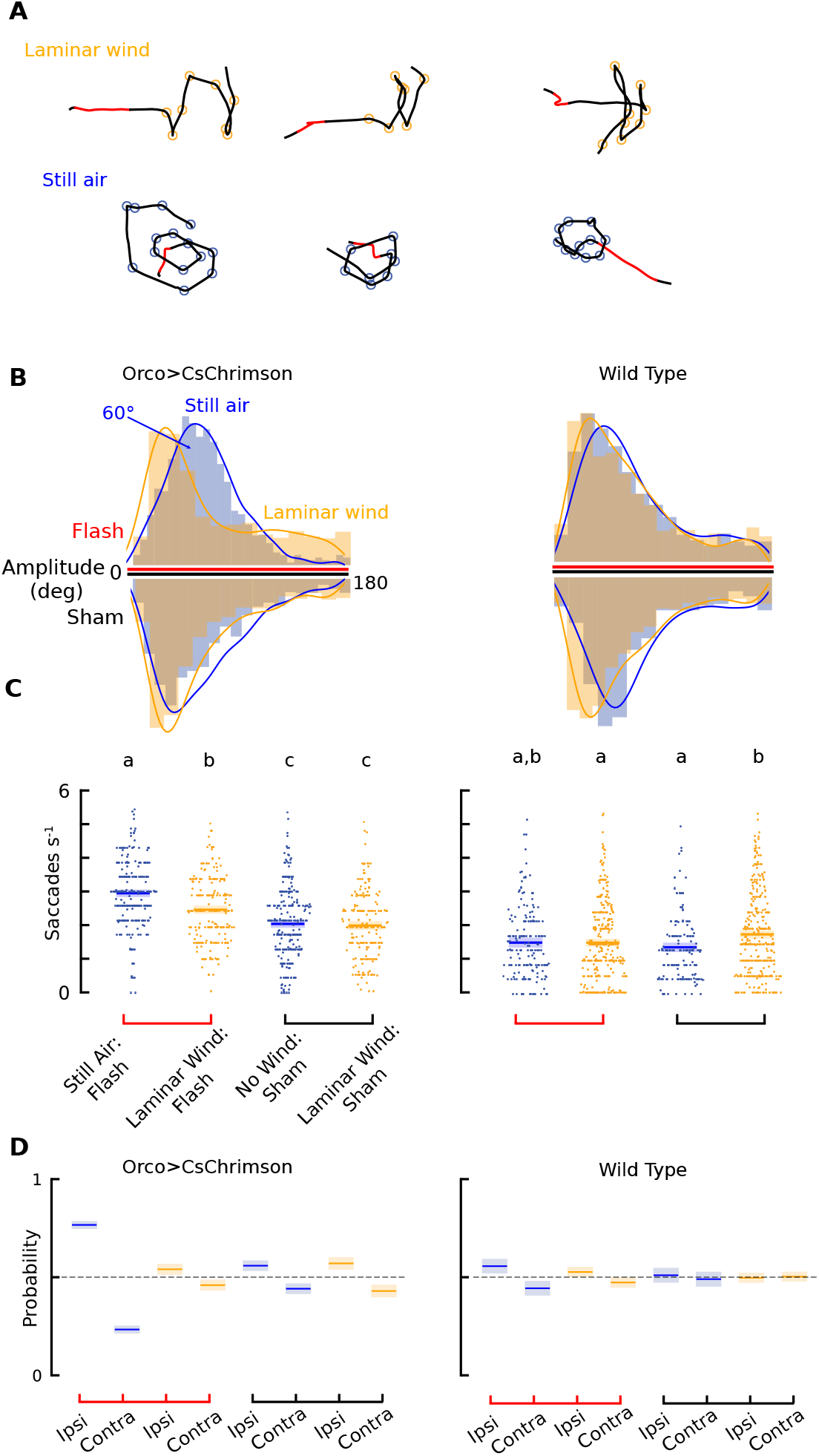
Saccade dynamics after optogenetic stimulus change in amplitude, frequency, and directional consistency based on the wind environment. (A) Six example trajectories of Orco>CsChrimson flies from either casting or circling conditions and where the mGSD algorithm annotated saccades (orange circles from laminar wind trajectories, blue circles from still air trajectories). (B) Saccade amplitude distributions from Orco>CsChrimson flies (left) in laminar wind (orange n=1230 flash, n=922, sham) or still air (blue n=1930 flash saccades; sham, n=1459). Distributions above the axis indicate trajectories from a flash event, and distributions below the axes indicate data from sham assignments. Wild-type controls are on the right hand side of the figure for B-D (laminar wind flash saccades, n=1134; laminar wind sham, n=1344; still air flash, n=695; still air sham n=648). (C) Saccade frequencies of trajectories from laminar wind or still air and their associated sham controls. Letters indicate statistical groupings within a genotype (Mann-Whitney U test with Bonferroni Corrected *α* = .0083). Striated tiers of points are from trajectories with full 3 second of post stimulus tracking, other points are from trajectories that had between 1 and 3 seconds tracking. Raw data points are distributed horizontally according to a histogram of the raw data, similar to a violin plot. (D) Ipsilateral vs. contralateral bias of saccades for each wind, flash, and genotype condition. We included only saccades from 0.9s-3.0 s after the stimulus onset to compare dynamics specifically from casting or circling behaviors. Lines on graphs represent means and error shading on graphical representations of data represent 95% confidence intervals. See also Figure S5.

In still air, Orco>CsChrimson flies’ post-stimulus circling saccades were a unimodal distribution around 60° (Figure 3B, mean = 59.1°± 31.2°). In contrast, Orco>CsChrimson flies engaged in post flash casting had a long tailed distribution of saccade amplitudes, mixed between frequent small saccades and less frequent large amplitude ones. The amplitudes of saccades from Orco>CsChrimson flies which received sham triggers in still air, wild-type flies in laminar wind and wild-type flies in still air all had similar long-tailed distributions, although casting flies used larger amplitude saccades more frequently (Figure 3B).

In addition to changes in the distribution of saccade amplitudes, Orco>CsChrimson flies which received a flash event in still air had a higher saccade frequency than those casting in laminar wind (still air, flash: 3.4 ±.95 Hz; laminar wind, flash: 2.9 ±.96 Hz) (Figure 3C). Orco>CsChrimson trajectories which were assigned sham events had similar saccade frequencies regardless of whether or not they were in still air or laminar wind (still air, sham: 2.5 ±1.0 Hz; laminar wind, sham: 2.4 ±1.0 Hz). Wild-type flies had lower saccade frequencies than Orco>CsChrimson flies, including Orco>CsChrimson flies which received sham events. We attribute this as likely being due to Orco>CsChrimson having some potential level of olfactory sensory neuron activation from white lights, and Orco>CsChrimsons flies’ prior experiences with red light flashes activating olfactory sensory neurons throughout the night. Both have the potential to change Orco>CsChrimson flies’ baseline behaviors compared to those of wild-type flies. Wild-type flies typically had similar saccade frequencies regardless of which trigger event they received, and whether or not they were flying in laminar wind or still air (still air, flash: 1.9 ±1.0 Hz; laminar wind, flash: 1.9 ±1.1 Hz; still air, sham: 1.8 ±.95 Hz; laminar wind, sham: 2.1 ± 1.2 Hz).

In still air, flies’ saccade dynamics became directionally consistent: they were nearly 3 times as likely to perform a saccade in the same direction as their prior saccade as opposed to the other direction (Figure 3D). This was significantly more ipsilaterally biased than any other condition (Z-test of proportions, Z=18.1, p=9.3×10^−38^). All other experimental conditions were either not ipsilaterally biased, or were biased drastically less so (Figure 3D). We found no obvious causality for whether a trajectory adopted a clockwise or counterclockwise bias.

### Circling is specifically in response to the loss of an odor

We next tested whether circling in still air was triggered specifically by the loss of the odor stimulus by implementing a “double flash” experiment. In these experiments, flies received either two 675 ms light flashes 1 second apart, or two shams (Figure 4A-B). We restricted our analysis to trajectories that extended for the full 3.4 seconds after the first stimulus. We pooled sham trajectories from the single and double flash experiments with the single flash experiments described earlier to make up for the fact that many sham trajectories were shorter than 3.4 seconds.

**Figure 4.**
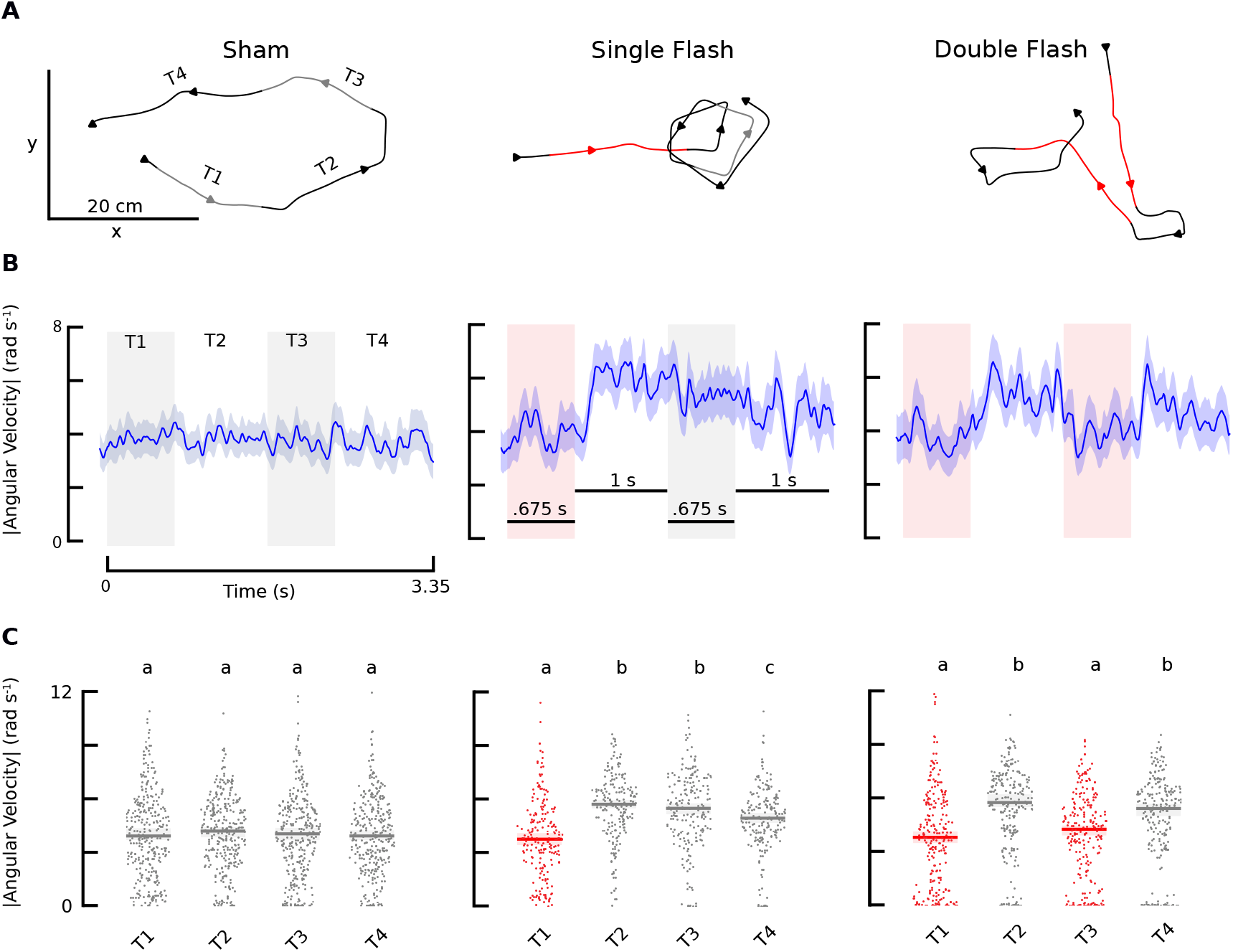
Circling is a response to odor loss. (A) 3.4 second sample trajectories in the x, y plane of flies that received no optogenetic stimulus, a single red light flash stimulus, or two flashes separated by one second. Red color indicates the portion of the trajectories during which the red light stimulus was active and gray indicates the portion of the trajectories which are temporally aligned with the optogenetic stimulus of the two flash experiment, but did not receive a red light stimulus during that time. (B) Mean population angular velocity responses for the respective no flash (N=165 flies: n=316 trajectories), single flash (N=60 flies: n= 200 trajectories), and double flash (N=105 flies: n=241 trajectories). (C) Mean angular velocities of individual trajectories over each of the four time regions of interest during their flight relative to entering the trigger zone: the first 675 ms (*T1*) the following intermittent second (*T2*), a secondary 675 ms portion (*T3*), and the subsequent one second period (*T4*). Lines on graphs represent means and error shading on graphical representations of data represent 95% confidence intervals. Letters represent statistical groupings that can be compared within a stimulus regimen. Statistical groupings were determined by Mann-Whitney U tests with a Bonferroni corrected *α* of .0083. Raw data points are distributed horizontally according to a histogram of the raw data, similar to a violin plot. See also Figure S4.

Sham trajectories showed no significant changes in mean angular velocity over the 3.4 seconds we analyzed. Flies receiving a single flash showed a sharp increase in mean angular velocity immediately after the stimulus ended (Figure 4B-C). Their angular velocity remained elevated for 2 seconds before tapering off slightly (Figure 4B-C). When flies received a second flash, they reduced their angular velocity during the stimulus, and subsequently re-initiated circling when the second flash ended (Figure 4B-C). We found no evidence of a visually mediated response to light flashes in wild-type controls (Figure S6).

### Search in still air is a proximal search strategy

We hypothesized that the behavior flies use in still air is likely a proximal search strategy. To show this visually, we aligned each trajectory in the horizontal plane around the point at which the fictive odor ended (Figure 5A-B). In still air, Orco>CsChrimson flies which received an optogenetic stimulus stayed significantly closer to the point at which the odor was lost compared to Orco>CsChrimson flies in laminar wind (Figure 5C).

**Figure 5.**
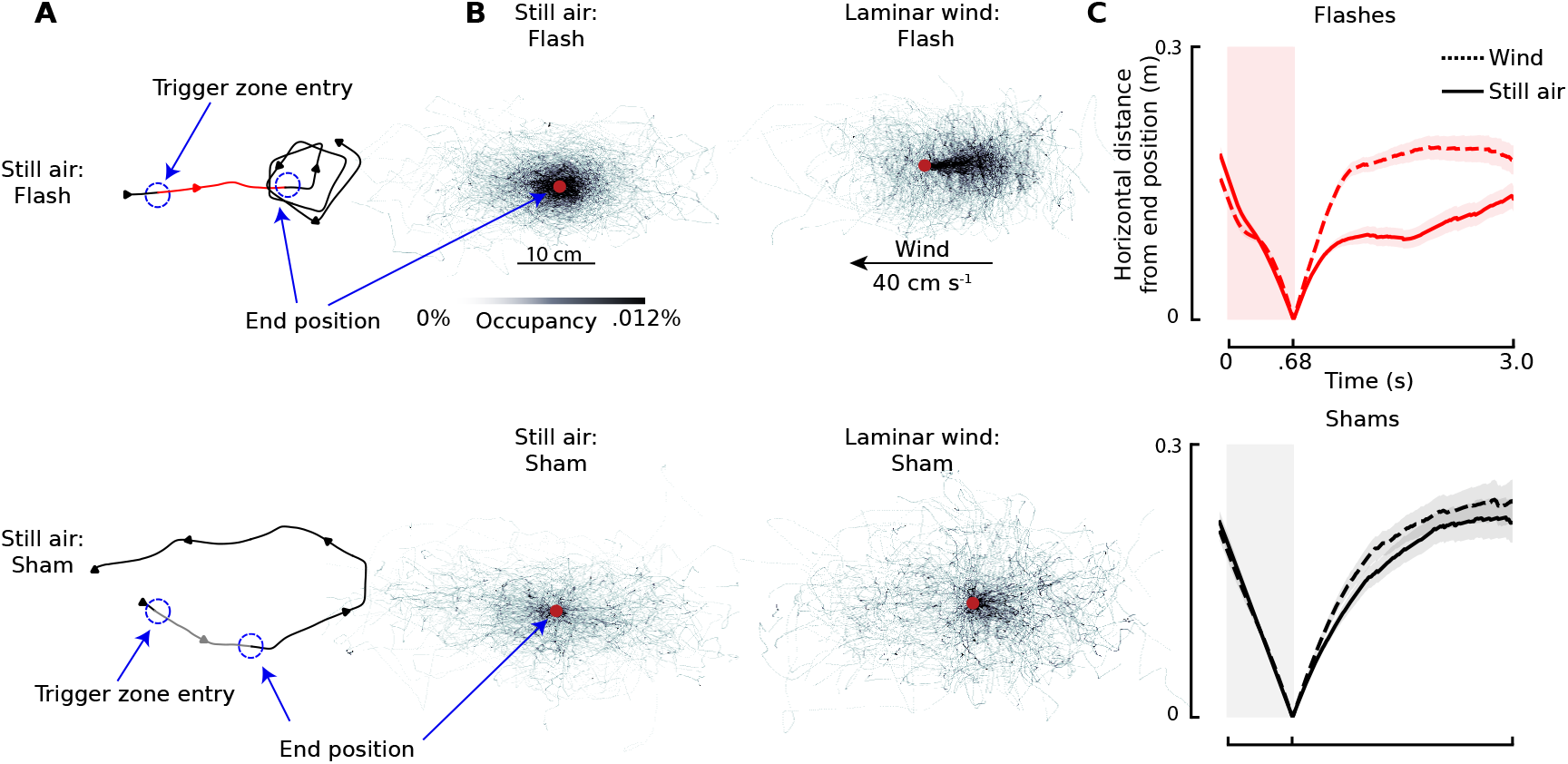
Search in still air is a proximal search strategy. (A) Example trajectories from flies in still air that received a flash (top) or sham (bottom) assignment. Red indicates portions of the trajectory during which the red light stimulus was active and gray indicates the 675 ms region after the sham trajectory entered the trigger zone. (B) Heat maps of trajectories with origin set to the flies’ position 675 ms after entering the trigger zone (i.e. when the stimulus turned off for flash events). Heat maps exclude the portions of the trajectories up until 675 ms after entering the trigger zone. Red dots indicate the point of alignment for all trajectories (i.e. when the stimulus turned off). (C) Mean horizontal distance from the end position for Orco>CsChrimson flies which received either a flash (top) (laminar wind: n=232 trajectories; still air: n=263 trajectories) or sham (bottom) (laminar wind: n=222 trajectories; still air: n=291 trajectories) assignment. Dashed lines are the population mean of flies from laminar wind experiments and solid lines represent flies from still air experiments. Lines on graphs represent means and error shading on graphical representations of data represent 95% confidence intervals. See also Figure S4.

This difference was greatest 1.1s after the end of the optogenetic stimulus; in still air flies stayed on average 10 cm closer to the point of odor loss compared to flies in laminar wind, the maximum difference was only 3.5 cm when comparing sham trajectories from both wind conditions. Wild-type flies did not travel as far in still air as in laminar wind, however this was explained by differences in ground speeds in the two wind conditions. The flash events had no effect on the distance they travelled (Figure S4).

#### Anemotaxis is often a rapid, two-turn sequence

Whereas the existence of wind gated search strategies suggests flies can estimate wind magnitude while flying, it is less obvious how they might do so. We suspect that flies estimate properties of the wind in the first few hundred milliseconds after activation of olfactory sensory neurons, during which time the behavior of flies is remarkably similar despite flying in still air or laminar wind. We found that in both laminar wind and still air, activating *Orco*+ neurons caused flies to decelerate (Figure 6A). For a fly flying in laminar wind, this deceleration event was followed by a sharp change in course direction. Flies flying in still air decelerated to a similar degree but had a more subtle course change response (Figure 6A). Neither wild-type flies nor genetic controls exhibited a deceleration or change in course in response to the red light flash (Figure 6A, Figure S4), indicating that both maneuvers are likely olfactory responses. In laminar wind, anemotaxis manifests as a second sharp turn that directly followed the initial course change, thus creating a sequence of two rapid turns (Figure 6B).

**Figure 6.**
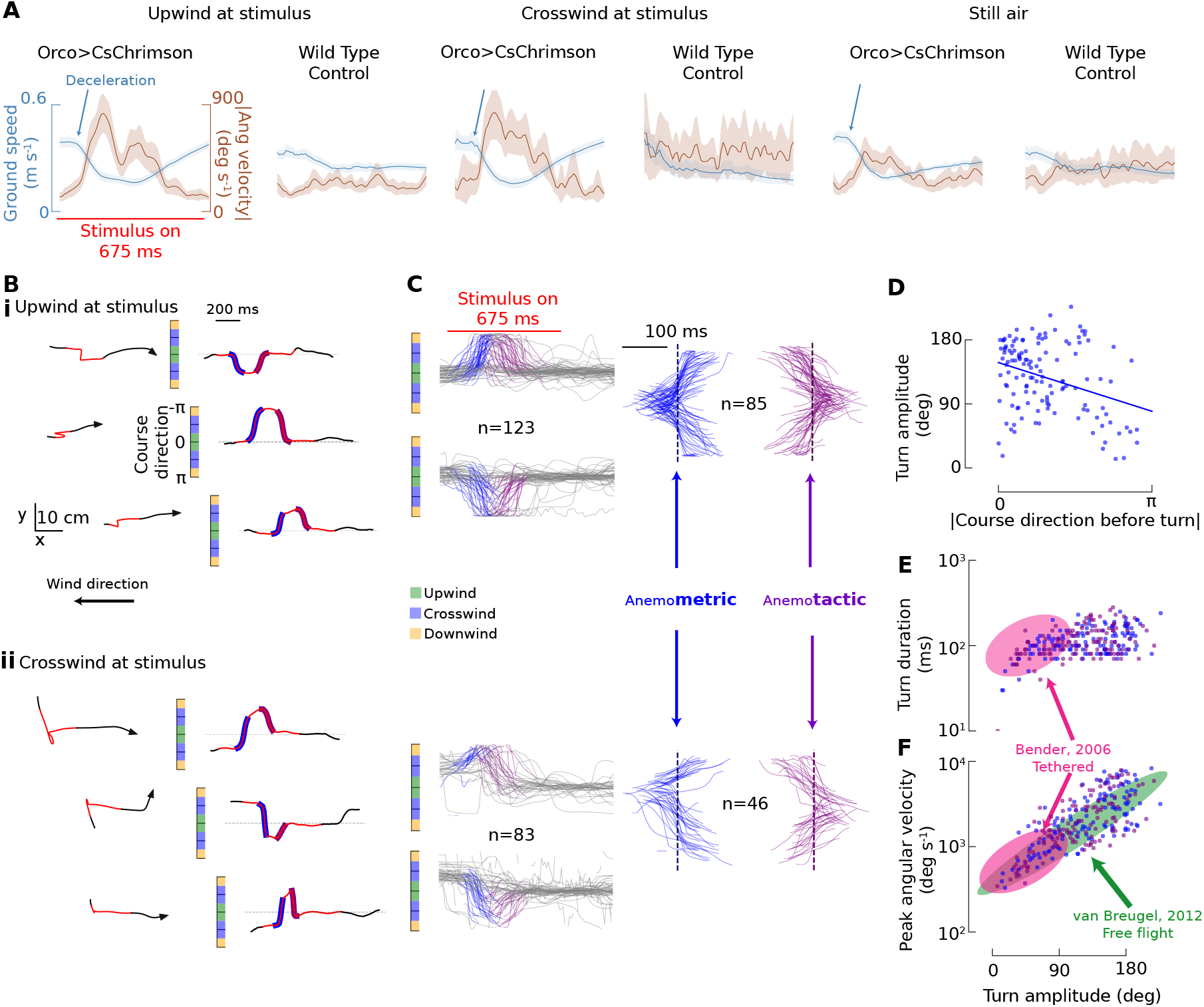
Anemotaxis is often a rapid two-turn sequence. (A) Population ground speed and angular velocity profiles for flies which received a flash of red light when entering the trigger zone. Groups were split into flies which were upwind at the time of the stimulus, crosswind at the stimulus, or in still air. (B) 1.1 second portions of trajectories as they orient into the wind and their associated course direction as a function of time for flies oriented either (i) upwind at the time of stimulus or (ii) crosswind at the stimulus. Course direction plots are highlighted blue for the initial (“anemometric”) turn and purple for the wind orienting (“anemotactic”) turn. (C) Course direction vs. time of flies as they orient into the wind, with the two turns extracted and centered around their midpoint to the right. Each anemometric turn has a corresponding anemotactic turn in this analysis. For the crosswind subset the sign of the course direction for some trajectories was modified to better visualize the turn regions. (D) Amplitude of the anemometric turn as a function of the trajectories course at the time of the stimulus. (E) Duration of the anemometric and anemotactic turns as a function of the turn amplitude. Blue markers are for the anemometric turns and purple markers indicate the anemotactic turns. (F) Peak angular velocity of turns as a function turn amplitude for the anemotactic and anemometric turns presented in this study. In (E-F) shaded ellipses show the approximate kinematic envelopes for body saccades exhibited by tethered *Drosophila*^16^ (pink), and rapid turns made by free flying *Drosophila*^29^ (green). Lines on graphs represent means and error shading on graphical representations of data represent 95% confidence intervals. See also Figure S6 and S7.

The two-turn anemotaxis sequence was most stereotyped in flies broadly oriented upwind at the time of the stimulus (n=123 trajectories, Figure 6C). Of the flies oriented crosswind at the time of the stimulus (n=83, Figure 6C) more than half also used a two-turn sequence to perform anemotaxis even often passing through an upwind heading during the initial turn before correcting with a second (Figure 6B-C). Flies oriented downwind at the time of the stimulus did not display stereotyped wind orienting behavior (n=26 total trajectories, Figure S6) and were not included in this analysis. The magnitude of the initial turn tended to be larger for flies oriented more upwind at the time of the stimulus (Figure 6D). Although not visually mediated, we cannot entirely rule out that this behavior may be linked to the somewhat unnatural odor experience of activating all *Orco*+ neurons simultaneously. However, we also found evidence that flies perform a similar two-turn sequence upon entering an ethanol plume using previously published data (Figure S7). Although fewer flies performed the two-turn sequence in response to the ethanol plume (37% compared to 69%), those that did used a similar kinematic profile. These results suggest that the two-turn sequence is an important, but previously overlooked component of olfactory navigation in flies.

We next asked if the two turns exhibit similar kinematics to body saccades, which are thought to be feed-forward motor programs which flying *Drosophila* use to execute rapid changes in course direction. We quantified the kinematics of both the initial turn, hereafter referred to as the putative “anemometric” turn, and the wind orienting turn, referred to as the “anemotactic” turn. Comparing the turns along two parameters: turn duration and peak angular velocity, both as a function of turn magnitude, revealed that both turns have kinematic profiles closely matching those of other rapid free flight turns and body saccades in tethered flying *Drosophila* (Figure 6E,F). Compared to voluntary body saccades in a tethered preparation^16^, flies in our experiment often made larger changes in course, yet the duration of these large turns remained close to 100 ms.

## DISCUSSION

### Summary

Here we establish that, like in larval and walking flies^32–37^, optogenetic stimulation of *Orco*+ neurons induces naturalistic olfactory navigation behaviors in free-flight. We then investigated how flies use information from their wind environment to deploy a plume tracking strategy. In still air flies use a previously uncharacterized plume tracking behavior dominated primarily by two stereotyped motifs: rapid saccades biased in a consistent direction while descending in altitude. In the same way that the casting element of “cast and surge” is a fly’s attempt to reconnect with a plume, we demonstrate that the circling element of “sink and circle” is a fly’s attempt to reconnect with an odor plume. The sinking element may facilitate sampling near the boundary layer. Under no slip boundary layer conditions, an odor is carried further downwind at a higher altitude and detection of the plume closer to the ground is likely only very near the source. It has been suggested that canids and murids alternate between bouts of rearing their snouts high in the air and lowering them close to the ground, as a strategy that facilitates odor source location^38,39^. Some fish species also modulate their position within the water column while searching for odors^40^. Insects may use altitude control as a convergent evolutionary strategy when tracking plumes. We hypothesize that sink and circle comprises a proximal search strategy. Detecting an odor packet in the presence of wind may not necessarily be indicative of proximity to an odor source as an odor packet can travel over large distances when carried by wind. However, detecting an odor in still air may indicate a high likelihood that a fly is close to the source and it is better to search locally. The approach we develop here may also be used in future studies to elucidate olfaction driven search in other wind scenarios, such as unsteady flow.

### Implications of circling and candidate supporting circuits

Evidence of circling behaviors can be found in prior modelling and behavioral studies. For example, trajectories of agents using an infotactic search strategy in still air superficially resemble circling flies^41^. Flying flies searching for an invisible odor in a cylindrical arena also resemble the searching behavior we describe here^27,42^ and walking flies search with high angular velocities after optogenetic activation of olfactory sensory neurons^43,44^. However, with a real odor, flies can dynamically interact with an odor plume and alternate between bouts of high turn rates while circling interspersed with stretches of straight flight after encountering an odor packet. This has the potential to generate the long-tailed Lévy distributions which have been previously associated with scale-free search in free-flying *Drosophila*^45^. Paradigms which can open the behavioral loop for plume tracking are therefore critical for understanding how models of complex search ethology can arise from the building blocks of simpler stereotyped motor programs.

Where in the brain are the motor control commands generated that result in the circling behavior? In the silk moth *Bombyx mori*, crosswind counter turning is likely generated by pairs ‘flip flop’ neurons in the lateral accessory lobe^46^. Saccade directionality and timing in flying flies may be controlled by analogous pairs of neurons^47^. Compared to walking moths however, crosswind casting turns in flies are not simply large alternating contralateral saccades, but are composed of a more nuanced structure of smaller saccades that together result in crosswind movement (Figure 3D). The stereotyped, rhythmic and unidirectional turns we describe here may prove to be an ideal candidate for investigating the neural mechanisms underlying long sequences of sensorimotor control.

### Anemotactic saccades imply flies possess an internal estimate of wind direction

Our observation that flies orient into the wind using a turn with saccade-like kinematics has important implications for understanding wind guided navigation. Saccades are feed-forward motor commands^48^ during which feedback from motion vision is actively suppressed^17,49^. This suggests that flies must have an internal estimate of the ambient wind direction in order to generate the feed-forward motor commands that drive their anemotactic turns. The rapid upwind turns, and the wind-gated decision to circle or surge, indicate that flies have an internal model for both the presence and direction of ambient wind. How flies calculate such an estimate and for how long it persists remains unclear. The behavioral time course in both laminar wind and still air is remarkably similar for the first ∼150 ms after odor onset, during which time flies decelerate. In the presence of wind, this deceleration event was often coupled with a saccade, and in still air flies decelerated to a similar degree but were less likely to perform a rapid turn. The deceleration event and the saccade event may be a set of active sensing routines to gauge properties of the wind while flying.

### Putative computational mechanisms for active sensing of wind presence and direction

For a fly flying on a straight course, it would be impossible to determine wind presence or direction. This is especially clear in the most extreme case–a fly moving directly upwind or flying straight in still air–where the sensory experience for both scenarios will be nearly identical. Recent control theoretic analyses have shown that if a fly were to decelerate or change course while maintaining altitude, however, the ways in which the magnitudes and angles of visual motion and apparent airflow change relative to one another can resolve key properties of ambient wind^19,22,50^. This section provides a brief review of these theoretical predictions for how insects might estimate wind direction, as well as new theoretical predictions for how insects might estimate wind presence, and finally we relate these predictions to our experimental results. Since flies maintain relatively constant altitude during their anemometric saccade and deceleration maneuvers (Figures 1E-F, 2D-E), we restrict our model to 2-dimensions, that is, we only consider sensor information and wind in a plane parallel to the ground. We also restrict our model to the assumption that wind remains constant across the time of the anemometric turn.

To estimate wind direction generally requires a change in course, and can be achieved using solely angular sensory measurements that are known to be encoded in the central complex. Figure 7A provides a graphical summary of our established active anemosensing hypothesis^22^ for wind direction estimation, which involves comparing the angles of perceived airflow and optic flow from either side of a turn (alternative solutions exist as well, for example, exchanging the angle of optic flow for the angle of linear acceleration^19^). The Law of Sines for the “wind triangle” before and after the turn, as well as its time derivative, yields three equations that contain three unique unknowns: the wind direction (*ζ*, which we assume is constant throughout the turn), and a normalized speed (ground speed relative to wind speed: *v* = *g*/*w*) before and after the turn. Provided that the fly changes its course (i.e. makes a turn), these three equations are independent and there is a unique solution. In other words, if the fly turns, there is a one-to-one mapping from the sequence of angular sensory measurements to the ambient wind direction^19^. It is unclear if, and how, flies might solve these equations, but it is mathematically feasible^22^. For interpretability, we describe the mathematics as a comparison between two points in time. In practice, it is more robust to implement a continuous comparison over time, which leads to greater robustness in the case of changes in wind direction^22^.

**Figure 7.**
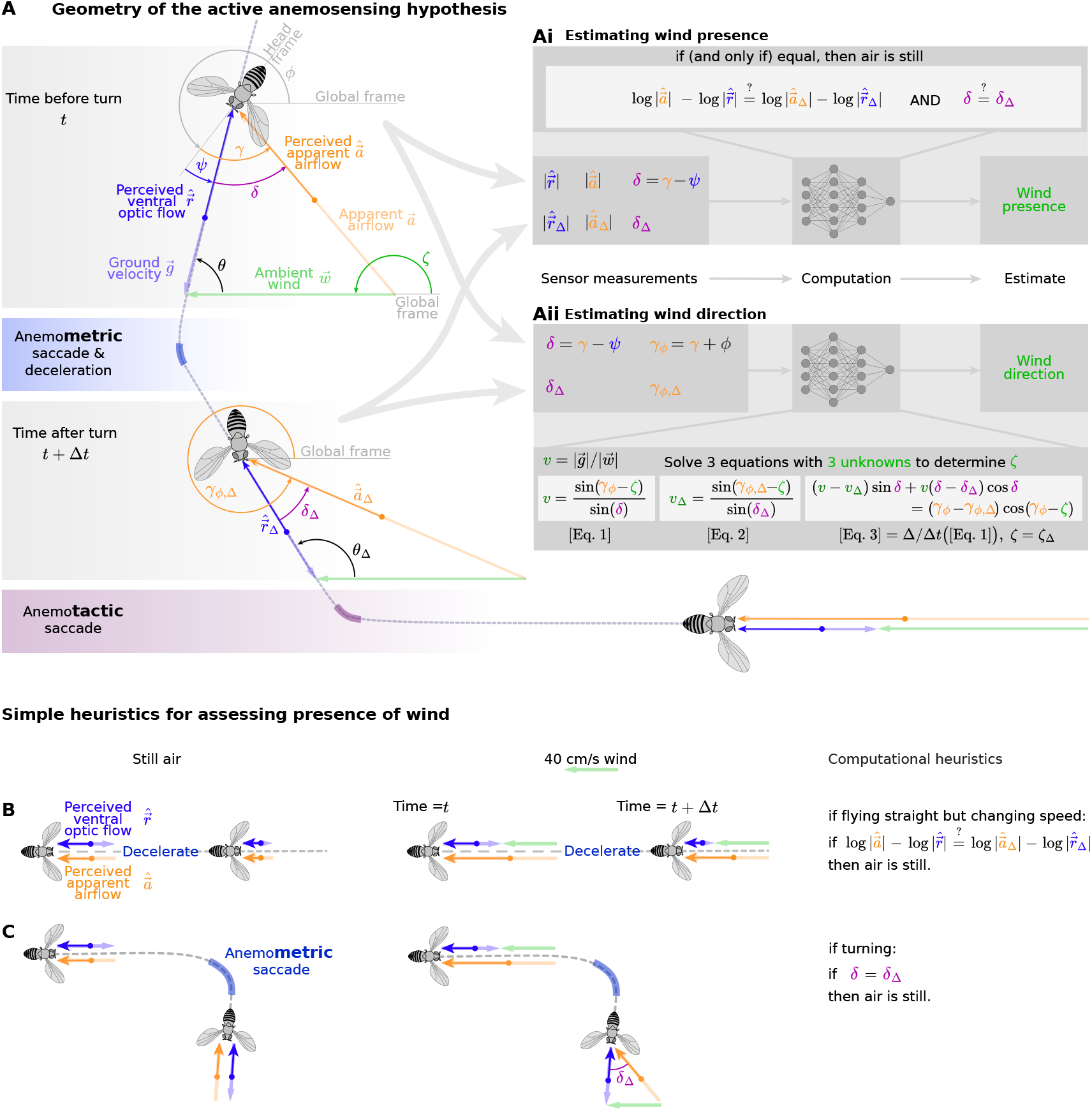
Review of active anemosensing hypothesis and candidate heuristic models for estimating presence of wind. (A) Geometry of the “wind triangle” before and after a putative anemometric saccade, followed by an open-loop anemotactic saccade. By comparing sensor measurements of perceived apparent airflow and optic flow from before (time=*t*) and after (time=*t* + Δ*t*) the anemometric turn, it is possible to estimate both wind presence and direction. (Ai) Proposed model of information flow for estimating wind presence, and a candidate mathematical description of the computations flies might approximate in their central complex to estimate presence of wind (see STAR methods for derivation). (Aii) Proposed model of information flow for estimating wind direction, and a candidate mathematical description of the computations flies might approximate in their central complex to estimate wind direction. Equations 1-2 correspond to the law of Sines for the two wind triangles. Equation 3 is a discretized time-derivative of Equation 1, assuming a constant wind direction of the time Δ*t* (see^22^ for derivation). (B) In still air, if a fly decelerates while maintaining course, the relative differences between perceived optic flow and apparent airflow magnitude remain constant (and vice versa). (C) If in still air a fly makes a saccade while flying at a constant ground speed, then the angle between its perceived optic flow and apparent airflow will remain constant (and vice versa). Note that the magnitude of the optic flow vector 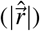 is both a function of ground speed and altitude, thus its instantaneous magnitude compared to the perceived apparent airflow magnitude does not provide unique information about the presence of wind; only the relative changes during deceleration would uniquely reveal the presence of wind.

To estimate wind speed, on the other hand, requires some type of magnitude information (without it, the scale of the “wind triangle” in Figure 7A cannot be determined). Broadly, changes in ground speed may facilitate estimates of wind speed. However, such estimates would require that flies have a stable calibrated measure of the magnitude of optic flow (see STAR methods). We can model a flies perceived ventral optic flow as 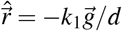, where 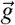 is the ground velocity vector, *d* is the altitude above the ground, and *k*_1_ is a calibration coefficient that includes effects such as the conversion of radians/sec to pixels/sec, as well as other effects such as the influence of contrast, motion adaptation, and state dependent modulation effects. This presents a complication because the responses of wide field optic flow sensitive neurons depend on texture and are subject to motion adaptation^51^, so *k*_1_ is unlikely to be stable over longer periods of time. Similarly, a fly’s perceived apparent airflow is also most likely scaled by some unknown coefficient: 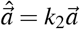. To address the challenges raised by unknown sensor calibration coefficients, we derived a binary heuristic for assessing the presence of wind across a combined deceleration and turning maneuver (Figure 7A, see STAR methods for derivation).

To provide better insight into our proposed heuristic for determining wind presence, we break it down into two delay and compare heuristics. First, consider a fly flying on a straight course. The angles of its perceived apparent airflow 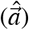 and perceived optic flow 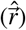 will be equal (i.e. *δ* = *γ* − *ψ* = 0) if the air is still, or the if the fly is moving straight upwind (or downwind very fast). In a *δ* = 0 scenario, the fly has two options to determine the presence of wind. It could stay on course and decelerate (Figure 7B) and compare the relative changes in magnitude of its perceived apparent airflow and optic flow. If 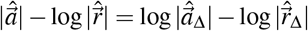 (and fly is moving straight, and therefore *δ* = *δ*_Δ_), then the air must be still, where 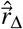 is a measurement of perceived optic flow from time Δ*t* after the measurement 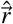 (see STAR methods for derivation). Alternatively, the fly could make a change in course while flying at a constant nonzero ground speed. In this case, if the relative angle between the perceived optic flow and airflow (*δ* = *γ* − *ψ*) remains constant before and after the turn, then the air must be still (Figure 7C, see STAR methods for derivation). In principle that relative angle *δ* should be zero both before and after the turn in still air. Comparing the relative values of *δ* to each other, instead of zero, makes this heuristic more robust to potential angular offsets in the sensory measurements.

Our experimental observations that flies decelerate, and often make a turn, after encountering an odor are consistent with the theoretical predictions outlined above. Based on this theoretical foundation, we refer to the turn flies make prior to turning upwind as a putative “anemometric” saccade, since we hypothesize that they use this turn to measure properties of the wind. In both still air and in laminar wind flies decelerate to a similar degree, whereas flies are more likely to perform a change in course when there is wind. Thus, we suggest that although flies may use a change in course to simultaneously assess wind presence and direction, we cannot rule out the hypothesis that they use a deceleration maneuver to assess the presence of wind independent of their decision to change course. Whether flies use these maneuvers to determine the presence and direction of ambient wind, and whether they use the candidate computational mechanisms we describe, remain open questions. Since we also found evidence of both maneuvers in flies interacting with a real ethanol plume (Figure S7), the deceleration and anemometric turn are unlikely to be visual artifacts, or artifacts of the unnatural odor percept from activating all *Orco*+ neurons simultaneously. The maneuvers could, however, either serve some other purpose in olfactory search, or they could be a general olfactory “startle” response.

There is clear evidence that three directional sensory modalities that would enable wind direction estimation are encoded in flies’ central complex–the direction of optic flow^52,53^, heading relative to landmarks^54^ or the sky polarization angle^55^, and the direction of apparent airflow^56,57^. The central complex also supports the necessary architecture for angular integration^53^, memory^58^, vector computations^59–61^ and turning-synchronized computations^62^ that information-gathering during course changes would require. Downstream neurons that project heading and sensory information to the fan-shaped body (PFN type neurons) encode both visual^59^ and airflow^33^ information, making the fan-shaped body a likely candidate brain region for computing ambient wind properties. This hypothesis is further supported by recent work indicating that flies encode a goal-state in the fan-shaped body^63^.

In principle it would be mathematically feasible for flies to update an internal model of ambient wind direction after every turn they make. However, we did not find evidence for a decrease in likelihood of performing an anemometric turn if they had made a recent prior turn. Given the propensity of olfactory cues to modulate fly behaviors and sensory processing, it is plausible that novel olfactory stimuli might trigger a behavior such as a wind estimation reflex. This might also help to explain why fewer flies perform the anemometric turn in response to the real ethanol odor (Figure S7), as those flies are dynamically interacting with a plume and therefore odor encounters are less likely to be novel encounters. To determine which turns flies use to estimate wind direction, and whether these calculations are modulated by olfaction, will likely require investigating neural activity during olfaction driven tethered flight behaviors. Again, the fan-shaped body is a likely brain region to start with since olfactory signals appear in fan-shaped body inputs^33^, and anatomical studies indicate that many fan-shaped body tangential inputs are downstream of the mushroom body^64,65^.

### Flow estimation across taxa

Understanding goal oriented navigation is a central objective of behavioral neuroscience. From a comparative perspective it is not well understood what sensory cues and internal states trigger localized search over long distance navigation. In flies we demonstrate that wind plays central role in the behavioral switch to highly localized search, but how flies, and organisms generally estimate properties of flow in their environment is poorly understood. Flow estimation is likely a general ability across organisms. Although the underlying behaviors and neural mechanisms remain poorly understood, fish exhibit similarly convergent behaviors and computations to flies. For example, lampreys will execute circling behaviors when they encounter a river plume before swimming upstream^66^, a behavior that may be analogous to the anemometric turns we describe in flies. Zebra fish rely on temporal changes of apparent flow measurements to deduce flow direction during rheotaxis^67^. The temporal changes of apparent flow are an important component of the theoretical foundations for flying flies to estimate both presence and direction of ambient wind without magnitude calibrated sensory systems. More broadly, our hypothesis of how flies may estimate wind presence and direction through active turns adds to a growing literature describing how organisms can use motion of their sensors to estimate features of their environment that are not directly measurable^68–71^.

## ACKNOWLEDGEMENTS

We thank Dr. Kathy Nagel for sharing *Drosophila* lines and for providing critical feedback at several stages of this project. We also thank Dr. Dennis Mathew and Dr. Tom Kidd who provided invaluable assistance and feedback from the early stages of this work. Drs. Michael Dickinson, Noah Cowan, Jeff Riffell, Ryan Tung, Christina May, and Ben Cellini also provided comments, which greatly improved the manuscript. We also acknowledge Dr. Matthew Smear and two anonymous reviewers whose comments greatly improved this manuscript. Austin Lopez provided critical assistance with the wind tunnel, and Kaylee Jamison provided assistance with fly care and preparation. Finally, we thank all of the members of the van Breugel lab who contributed ideas and suggestions for this manuscript from its nascent stages. This work is supported by: NIH (P20GM103650); AFOSR (FA9550-21-0122); Sloan (FG-2020-13422); NSF-AI (2112085) all to FvB.

## AUTHOR CONTRIBUTIONS

Study conception and design: SDS and FvB. Data collection: SDS. Analysis and interpretation of results: SDS and FvB. Mathematical derivation: FvB. Manuscript preparation: SDS and FvB.

## DECLARATION OF INTERESTS

The authors declare no competing interests.

## STAR METHODS

### RESOURCE AVAILABILITY

#### Lead Contact

Further information and requests for resources and reagents should be directed to and will be fulfilled by the lead contact, Floris van Breugel

#### Materials Availability

This study did not generate new unique reagents.

#### Data and code availability

Data will be made publicly available on Dryad upon publication. All analysis code used in this paper is publicly available online at: https://github.com/dstupski/wind_gates_search_states.

### EXPERIMENTAL MODEL AND SUBJECT DETAILS

#### Drosophila melanogaster

All *Drosophila melanogaster* were maintained in a temperature (25°C) and humidity controlled incubator (60% RH) on a 16:8 light:dark cycle. All flies were reared on standard fly media (Nutrifly-Bloomington Formula, Genesee Scientific, San Diego, CA, USA). The F1 flies from crosses for experiments involving optogenetic stimulation were reared in a fresh media bottle pre-treated with 400 *µ*L of a 40 mM ATR solution (ATR, R2500, Sigma Aldrich) and maintained on the fresh ATR enriched media for at least 48 hours prior to experimentation. Experimental flies (2-5 days old) were anesthetized at 4 °C on a cold plate, and 15 females were placed in an empty tube with a moist Kimwipe to prevent desiccation. We then starved flies in their preparation tubes for 6-8 hours in the morning prior to any experimentation to encourage flight and food seeking. In the evening, we placed experimental flies into the wind tunnel behavioral system without any further anesthesia. Flies were permitted to fly about the tunnel volume overnight until the following morning at which point the data collection was terminated. All experiments included a minimum of two independent overnight recording bouts from 30 total flies-but usually four or more recording bouts from 60 flies. Although our study only used female flies, male flies likely respond in a similar manner. We used female flies because their larger size makes them more robust for longer experiments, and easier to track with our tracking system.

### METHOD DETAILS

#### Behavioral Arena

Our free flight behavioral arena consisted of a custom-built wind tunnel with a 1.0×0.5×0.5 m^3^ working section (Figure 1A), housed in a room maintained at 22° C and 60% RH. While experiments were running, we continuously ran a carbon air filtration system (SS-422, Sentry Air Systems, Cypress, TX, USA) to remove ambient odors. The airflow in the wind tunnel was generated by an array of computer fans controlled by an Arduino Mega 2560 board. For laminar wind experiments, we set the wind speed in the tunnel to 40 cm/s (calibrated using an ultrasonic anemometer: Trisonica Mini, Anemoment LLC / LiCor, Longmont, CO, USA). For still air experiments, all conditions were the same with the exception that the wind tunnel fans were not running, and we confirmed that there was no appreciable air flow within the wind tunnel using an anemometer.

For illumination within the tunnel, we outfitted each wall of the wind tunnel with arrays of blue LEDs (STN-BBLU-B6A-08C1M-24V, Super Bright LEDs Inc. St. Louis, MO, USA) and infrared LEDs (PN: 7031, Waveform Lighting, Vancouver, WA, USA). The side walls and bottom were lined with a light diffusing film (Spye Smoke, Spyeglass, Minneapolis, MN) to generate smooth infrared lighting over the wall surface (Figure 1A). Infrared LED’s lined the entirety of each wall to generate an imperceptible bright background to track flies against. The blue LED’s homogenously illuminated the wind tunnel floor, but were arranged along the wind tunnel walls to create an ombré effect that tapered from blue on the wind tunnel floor to black towards the wind tunnel roof. Along the wind tunnel floor we placed a grid of dark, infrared transmissible film, to create a checkerboard effect and provide flies with visual contrast. For experiments investigating attraction to a visual feature on the wind tunnel floor, we did not include the checkerboard lattice and instead placed one dark, infrared transmissible acrylic disk (3.8 cm diameter) on the wind tunnel floor. We complemented the blue and infrared illumination in the wind tunnel with an array of white LEDs that ran along either side of the top of the wind tunnel to provide sufficient orange wavelength light for photoreceptor reisomerization^72,73^. Total illumination of the combined white and blue lighting in the wind tunnel was 425 lux in the wind tunnel center, measured by a LX1330B light sensor (Dr. Meter, Hong Kong).

#### 3-D Tracking

Real time 3-D tracking of flies was performed at 100 hz using an array of 12 Basler cameras (acA720, Basler AG, Ahrensburg, Germany) positioned above the wind tunnel (Figure 1A). Cameras were equipped with IR-pass filters so that tracking was performed using only infrared light. 3-D tracking was implemented in the open source Braid software^25^. All cameras’ intrinsic warping parameters were calibrated using a back illuminated 5×7 inch checkerboard and the built-in checkerboard calibration functionality within Braid.

#### Optogenetic Stimulus and Experimental Paradigm

Experimental paradigms were developed using custom software in C++ and Python, and coordinated with the tracking software via the Robot Operating System (ROS).

Our optogenetic stimulus paradigm used the real time 3-D position of a fly to inform a triggering function which was activated when the fly crossed into a pre-defined “trigger zone” within the wind tunnel (Figure 1B). When a fly entered the trigger zone it was randomly assigned either a pulse of light, referred to as “flash” throughout, or no light pulse, referred to as “sham”. The optogenetic stimulus itself was generated by powering the red channel of two arrays of Triple Bright RGB LEDs (SparkFun Electronics, Niwot, CO, USA) that lined both the left and right-hand sides of the top of the wind tunnel (Figure 1A). Because the light environment was not entirely homogeneous we generated a 3-D model of the intensity of red light within the trigger zone of the wind tunnel, the average of which is our reported stimulus intensity, 42 *µ* W *mm*^−2^ (Figure S1). The delay between tracking software and activation of the optogenetic stimulus array was approximately 6 ms. Because 6 ms was less than the interval between individual camera frames, we considered the optogenetic stimulus delay to be negligible for the purposes of our analyses. We included random sham triggering events to account for the closed loop behavior of what a fly would naturally do in the wind tunnel volume in the absence of any kind of optogenetic perturbation. Comparing the flash and sham conditions for the wild-type allowed us to check for visually mediated artifacts. Comparing the flash and sham conditions for the experimental Orco>CsChrimson shows that the behaviors we observe are the result of olfactory receptor neuron activation above any baseline activation due to the white lights in the wind tunnel. For flies that triggered a light flash, the stimulus consisted of a 675 ms continuous pulse of light unless otherwise noted. We chose 675 ms because it clearly allowed us to separate cast and surge states for the majority of flies (Figure 1D).

Experiments included a 5 second refractory period during which flies could not reactivate a triggering event. Because trajectories could enter a far upwind or downwind zone within the wind tunnel where tracking was not possible, we were often limited to only a few seconds of behavior after each triggering event. We therefore restrict most analyses to the 3 second period directly after a flash or sham event.

#### Plume Generation

For the subset of experiments which involved an ethanol plume, we continuously pumped air through a 50% ethanol solution and up through a port that arose from the bottom of the wind tunnel on the far upwind end. We controlled the flow rate of the plume at 130 *cm*^3^ /minute using a digital mass flow control system (Alicat Scientific, Tuscon, AZ, USA). The port was connected to a 25 cm tall rigid acrylic tube that was bent at a 90°angle and with its open face covered in a metallic mesh to prevent flies from entering the tubing. This generated an ethanol plume in approximately the same volume as the optogenetic trigger zone.

#### Trajectory Inclusion Criteria and Data Filtering

For all optogenetic experiments, we included only trajectories which specifically elicited a triggering event (flash or sham). We also only included trajectories for which we had at least one full second of tracking after the triggering event before losing track of the fly when it entered an occluded zone within the wind tunnel. In the unlikely occasion that the same trajectory re-triggered the optogenetic stimulus we restricted analyses only to the first triggering instance for a given trajectory. On rare occasions, we found trajectories with brief jagged portions. We filtered these trajectories by excluding data points where a fly’s ground speed was greater than 2.0 m/s, an uncharacteristically high ground speed for *Drosophila melanogaster* flying either outdoors, or in a constrained space^74–76^. Additionally, because angular velocity measurements twice propagate the derivatives of tracking noise from flies’ x and y position, we found that angular velocity estimates on occasion experience brief highly variable segments along a trajectory. To account for this in our angular velocity analyses, we implemented a Butterworth Filter on the trajectory course direction before first order finite differentiation, all implemented in Python using the PyNumDiff package^77^.

#### Saccade Detection

Fly fruit flies typically change course via rapid turns referred to as body saccades^78^. To identify and characterize saccades, we used a slightly modified version of the Geometric Saccade Detection algorithm found in^30^. Our primary modification to the original algorithm was to include a term that took a trajectory’s ground speed into account in determining where saccades occurred (STAR methods). We visually inspected trajectories for agreement on saccade annotations (Figure S5). We did find that some trajectories included erroneous saccade labelling, likely arising from tracking noise that resulted from a fly leaving the field of view of one tracking camera or entering the field of view of another. We excluded these trajectories (¡15% of all trajectories) from analyses but the decision to include or remove them had no noteworthy impacts on conclusions drawn from the data. Kinematic properties of saccades were determined by isolating the portion of the trajectory 70 ms before and 70 ms after the saccade annotation.

#### A Note on Experimental Flies, Genetic Controls, and Characterizing Visually Mediated Artifacts

Motivated by prior optogenetic activation experiments with walking flies^32,43,44,79,80^, we chose to express the red-light sensitive ion channel, CsChrimson in olfactory receptor neurons that express the co-receptor *Orco* using the Gal4-UAS binary expression system. Although *Orco*+ neurons mediate a broad range of olfactory responses that are linked to odors with a range of valences^81,82^, the behaviors we observed by activating *Orco*+ neurons (detailed in results) are remarkably consistent with published free flight responses of *Drosophila* tracking ethanol and vinegar plumes in flight (Figure 1 and S2)^8^.

Unlike much of the prior work in walking flies, however, we could not use visually blinded flies because flight requires visual feedback. This raises the possibility of inducing visually mediated startle responses from sudden flashes of red light, despite flies’ relatively low red light sensitivity^83^. This is further complicated by standard genetic practices which use flies made in genetic backgrounds with a loss of function mutation in the *White* gene, the protein product of which is necessary for synthesizing the red screening pigments in the adult fly eyes. We therefore elected to use experimental flies which were at least heterozygous for a wild-type copy of *White* which resulted from crosses using a parental line that was homozygous for both the GAL-4 and UAS elements with wild-type flies (Key Resources Table). To characterize any visually mediated startle responses from red light flashes in genetic controls, we prioritized matching the *White* genotype of our Orco>CsChrimson flies (Figure S2). Our paradigm resulted in negligible visually mediated artifacts in two genetic controls: CsChrimson/+ flies and wild-type flies. We did find that our Orco>CsChrimson flies raised on media which had no ATR supplementation were moderately responsive to red light flashes. However, we attributed this as more likely due to weak olfactory sensory neuron activation rather than a visual response because they displayed a weak, time delayed anemotaxis behavior (Figure S2). After establishing that there were no salient visually mediated startle effects and due to the time consuming nature of our behavioral experiments, we elected to simply use wild-type flies as genetic controls for experiments in still air.

#### Modified Geometric Saccade Detection Algorithm (mGSD)

For saccade analyses and annotation of where saccade events occurred, we used a modified version of the Geometric Saccade Detection Algorithm found in Censi et al^30^ Supplemental Section 7.1. We calculated the mGSD score as follows.

1) For a given flight trajectory in the horizontal plane define a trajectory as *p*⟨*k*⟩ = ⟨*x*(*k*), *y*(*k*)⟩ where *x* and *y* are the x and y positions of the fly at frame *k*. Here *k* is an integer ranging from 1 (the first frame in the region of interest of a given trajectory) to n (the last frame in the region of interest along a trajectory).
2) Define a time step *δ*, in our case 5 frames (50ms), and for all *k* from 1+*δ* to n-*δ*). Redefine the origin of the coordinate frame so that 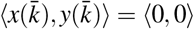.
3) For each 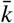 define two intervals, the “before” interval 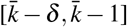 and an “after” interval 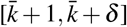.
4) For both the before and after intervals, calculate the angle, *θ* of each ⟨*x*(*k*), *y*(*k*)⟩ with respect to the new origin.
5) We define two angles, *θ*_*before*_ and *θ*_*after*_, as the median angle of the before and after intervals respectively. We then define an amplitude score at 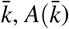, as:

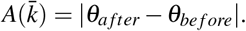
6) We then calculate a dispersion score, 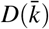 based on the horizontal distance travelled by a fly within +/-*δ* around 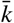. We define the dispersion parameter as the standard deviation of the flies linear distance from the newly defined origin over the 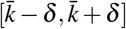 interval.
7) At 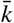 the mGSD score, 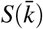, is then:

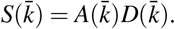
8) To determine at which frames a saccade event occurred we selected a threshold score for 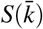 of 0.001. We then found all continuous regions over a trajectory where the saccade score was greater than our threshold, and during which it did not drop below the threshold for more than 5 frames in a row. We then annotated saccade events as the median position of the trajectory of each of these continuous runs where the score was over the threshold.

#### Volumetric Stimulus Calibration

We generated a 3-D model of the light intensity within the trigger zone volume by coupling the real time Braid tracking with light intensity measurements from an FDS100 photodiode (Thor Labs, Ann Arbor, MI, USA). We constructed a module that housed four infrared LED’s equidistant from the photodiode (Figure S1A). We inferred the photodiode position as the centroid of the four infrared LED positions, which were tracked using Braid (Figure S1B). We slowly moved the light measurement module throughout the volume of the wind tunnel with all of the room and wind tunnel lights turned off except for both of the arrays of red LED’s which lined the top of the wind tunnel. While moving the light measurement module we synchronously recorded the photodiode position and a voltage measurement from the photodiode itself (Figure S1C). We converted raw voltage measurements to light intensity measurements using the FDS100 data sheet and assuming a red light wavelength of 630 nm. From the raw position measurements we created a 3-D mesh of the light intensity within the volume of the pre-defined trigger zone (Figure S1D). Our reported stimulus is the mean light intensity in the trigger zone mesh, 42 *µ* W *mm*^−2^.

#### Derivation a Heuristic for Assessing the Presence of Wind

To determine wind speed, a fly would need to integrate information about its perceived airflow^1^ with an estimate of its ground velocity. Although insects cannot directly measure ground velocity, they do have access to the magnitude of ventral optic flow, which provides information about the ratio of ground velocity to their altitude above the ground. We can describe the ventral optic flow perceived by the fly as 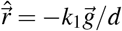, where 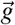 is the ground velocity, *d* is the altitude, *k*_1_ is a calibration coefficient that includes effects such as the conversion of radians/sec to pixels/sec, as well as other effects such as the influence of contrast, motion adaptation, and state dependent modulation effects. A simple solution for assessing the presence of wind would be for a fly to decelerate until it is hovering relative to its visual reference 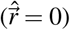. When hovering relative to a stationary visual reference the airflow magnitude will only be zero if there is no wind. However, we found that on average flies do not slow down to that extent. By accelerating or decelerating, it would be feasible for an insect to estimate its ground speed and the distance to objects^84,85^, and by extension, wind speed as well^50^. Such a calculation would, however, require that the magnitude of optic flow is accurate and calibrated–i.e. the value of *k*_1_ must be known. The responses of wide field optic flow sensitive neurons in flies are, however, dependent on spatial frequency and subject to motion adaptation^51^, indicating that for a fly the value of *k*_1_ is not stable over long periods of time, and its value may not be known. Meanwhile, although flies likely have access to a measurement correlated with airflow magnitude, it may not necessarily be calibrated with respect to the same units as their representation of optic flow. Thus, we include an unknown calibration coefficient in our model of flies’ perception of airflow as well. We describe the airflow perceived by the fly as 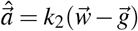, where 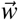 is the ambient wind vector.

To account for both unknown calibration coefficients (*k*_1,2_), we derive a simple delay and compare heuristic for assessing the presence of wind that is independent of these coefficients. After deriving a general two-dimensional heuristic, we consider the two special cases given in Figure 7B-C.

In some scenarios the presence (but not the absence) of wind might be trivial to determine from the trigonometry of the wind triangle (see Figure 7A). If the angle between the perceived airflow and perceived optic flow (*δ* = *γ* − *ψ*) is not zero, then there must be wind. If *δ* is zero (or very small), however, this could correspond to there either being no wind, or the fly moving directly upwind, or moving downwind faster than the wind is blowing. To resolve this ambiguity, flies could compare the relative changes in the vectors of their perceived airflow and optic flow across a deceleration and/or turning maneuver. To illustrate how this could work with more mathematical rigor, consider a scenario where a fly is maintaining a constant altitude in constant wind, or in still air (note that during the deceleration maneuvers we observe, flies do not, on average, change their altitude by very much (Figures 1E-F, 2D-E)). We write the following expressions for perceived ventral optic flow and air flow, taken from before and after a maneuver (or at any point during the maneuver):

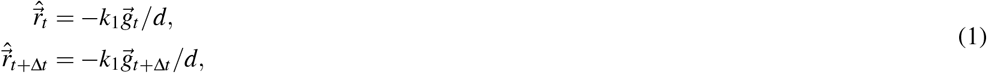

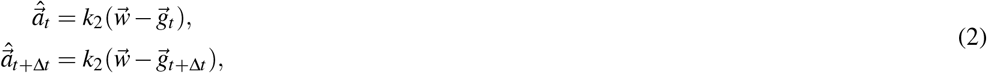

where *d* is the fly’s altitude, 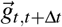 are the ground velocity vectors at two points along the trajectory, 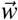 is the constant wind velocity vector, *k*_2_ is the calibration coefficient for airflow measurements, and *k*_1_ is the calibration coefficient for optic flow measurements. We assume that both *k*_1,2_ are constant over the duration of the maneuver. We assume the fly has access to 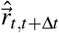 and 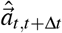, but none of the other variables. To simplify the expressions going forward, we introduce the following notation:

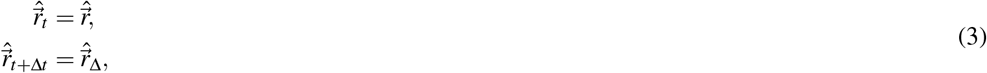

and similarly for other vectors and angles.

As discussed above, a key indicator of the presence of wind is the relationship between the perceived airflow and perceived optic flow vectors across time. Since the optic flow vector is aligned with the ground speed vector, we construct our heuristic by comparing the air and optic flow vectors with the ground speed vector using vector dot products (·):

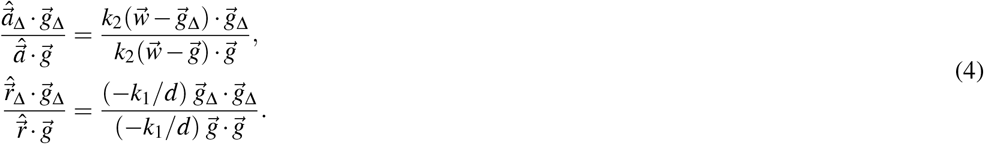

The unknown calibration coefficients on the right-hand side cancel, so we can simplify to:

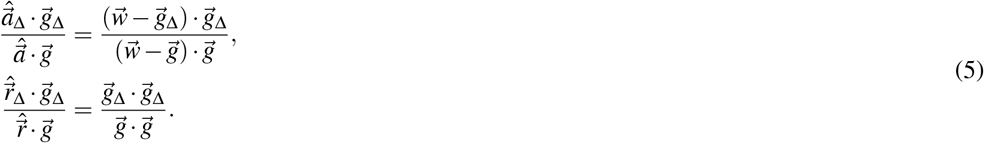

Evaluating the dot products on both sides of each equation yields the following:

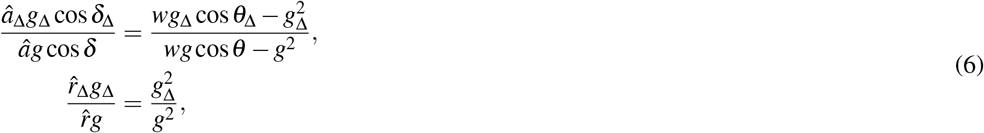

where 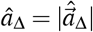, and similarly for the other vectors, and *θ* is the angle between the wind vector and ground velocity (as in Figure 7). Simplifying by dividing both sides of both equations by *g*_Δ_ and multiplying by *g* (and asserting that *g*≠0 and *g*_Δ_≠0) yields:

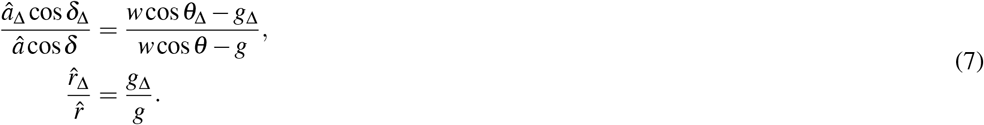

Now we ask under which conditions the right hand sides of these expressions are equal to each other:

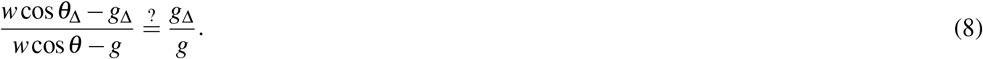

Multiplying the denominators, we get:

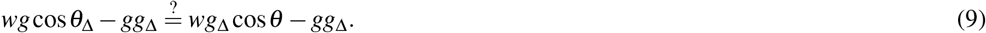

This equality could be true under any of the following four conditions:

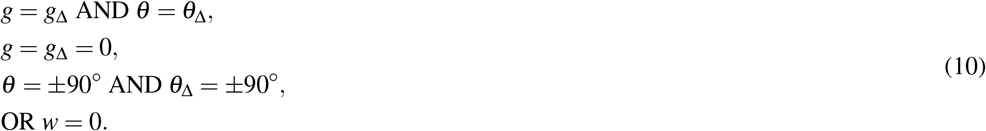

If we require that *g*≠0 and *g*_Δ_≠0 as established earlier, and that either *g*≠ *g*_Δ_ or *θ*≠*θ*_Δ_ (that is, either the fly changes its ground speed or turns relative to the ambient wind vector, which the fly would register as a change in *φ* + *ψ*), then we are left with only two conditions:

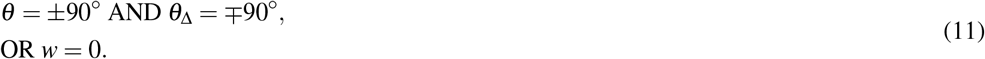

Thus, either the ambient wind speed must be zero, or the fly must have alternated between flying in the two directions perpendicular to the wind. Note, however, that if there is wind the latter case would correspond to relatively large values for *δ, δ*_Δ_, and their values would be equal in magnitude and opposite in direction. We established earlier that if *δ* or *δ*_Δ_ is not zero, then there must be wind. We can make the heuristic more robust to additive calibration offsets in the measurement of the angle *δ* by requiring that *δ* = *δ*_Δ_, but not necessarily requiring these angles to be zero.

Thus we can define a two part heuristic:

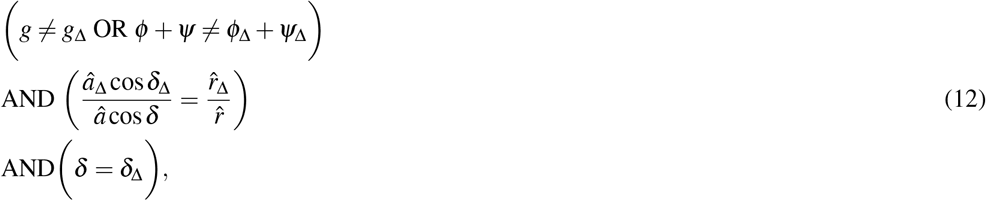

then the air is still.

Given that *δ* = *δ*_Δ_ for this heuristic, we can simplify to:

if, and only if:

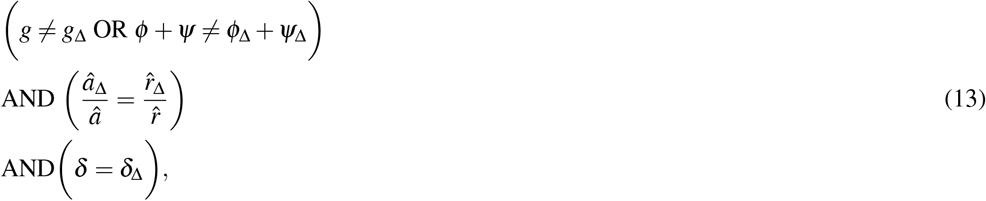

then the air is still.

To make it easier to see how such a heuristic might be implemented in a biologically plausible way, we can take the log of the first condition, arriving at two simple delay and correlate equality’s:

if, and only if:

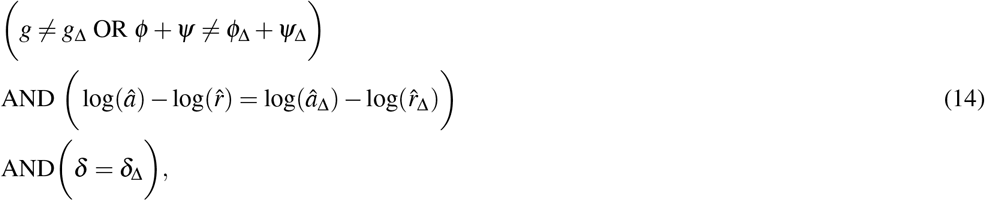

then the air is still.

To visually see the geometric implications of this heuristic, it is helpful to consider the two special cases shown in Figure 7B-C where either *g*≠*g*_Δ_ and *φ* + *ψ* = *φ*_Δ_ + *ψ*_Δ_ (Figure 7B), or *g* = *g*_Δ_ and *φ* + *ψ*≠ *φ*_Δ_ + *ψ*_Δ_ (Figure 7C).

### QUANTIFICATION AND STATISTICAL ANALYSIS

All statistical analyses were performed in Python 3.8 using SciPy Stats version 1.9.2^86^ or StatsModels^87^. Because we could not maintain fly identity over the course of an experiment, we assumed all trajectories are independent samples. Each experiment presented however, is the compilation of a minimum of 3, 16 hour recording periods from at least 45 independent flies. For all saccade specific analyses we considered saccades to be independent events, although we verified that if we instead averaged the properties of saccades from each individual trajectory and use each trajectory as an independent sample, the results are consistent. Data are reported in terms of means and standard deviations. Lines on graphs represent means and error shading on graphical representations of data represent 95% confidence intervals.

### KEY RESOURCES

| Experimental models |  |  |
| --- | --- | --- |
| Experimental Genotype | Parental Strain (s) | Source |
| NorpA/+;Orco-Gal4/+;UAS-CsChrimson/+ | NorpA;Orco-Gal4;UAS-CsChrimson and Heisenberg-CantonS | Dr. Katherine Nagel<br>Dr. Michael Dickinson |
| wild-type | Heisenberg-CantonS | Dr. Michael Dickinson |
| UAS-CsChrimson/+ | UAS-CsChrimson and Heisenberg-CantonS | BDSC_55135<br>Dr. Michael Dickinson |

## VIDEO LEGENDS

**Video S1. Representative 3-dimensional plume tracking trajectory in laminar wind. Related to Figure 1**.

**Video S2. Representative 3-dimensional plume tracking trajectory in still air. Related to Figure 2**.

**Video S3. Representative 2-dimensional plume tracking trajectories in still air. Related to Figure 2**.

**Video S4. Representative 2-dimensional plume tracking trajectories in still air. Related to Figure 2**.

**Video S5. Representative 2-dimensional plume tracking trajectories in still air. Related to Figure 2**.

**Video S6. Representative 2-dimensional plume tracking trajectories in still air. Related to Figure 2**.

**Video S7. Representative 2-dimensional plume tracking trajectories in still air. Related to Figure 2**.

**Video S8. Representative 2-dimensional plume tracking trajectories in still air. Related to Figure 2**.

**Video S9. Representative 2-dimensional plume tracking trajectories in still air. Related to Figure 2**.

**Video S10. Representative 2-dimensional plume tracking trajectories in still air. Related to Figure 2**.

**Video S11. Representative 2-dimensional plume tracking trajectories in still air. Related to Figure 2**.

**Video S12. Representative 2-dimensional plume tracking trajectories in still air. Related to Figure 2**.

**Video S13. Representative 2-dimensional plume tracking trajectories in still air. Related to Figure 2**.

**Video S14. Representative 2-dimensional plume tracking trajectories in still air. Related to Figure 2**.

## SUPPLEMENTAL MATERIALS

**Figure S1.**
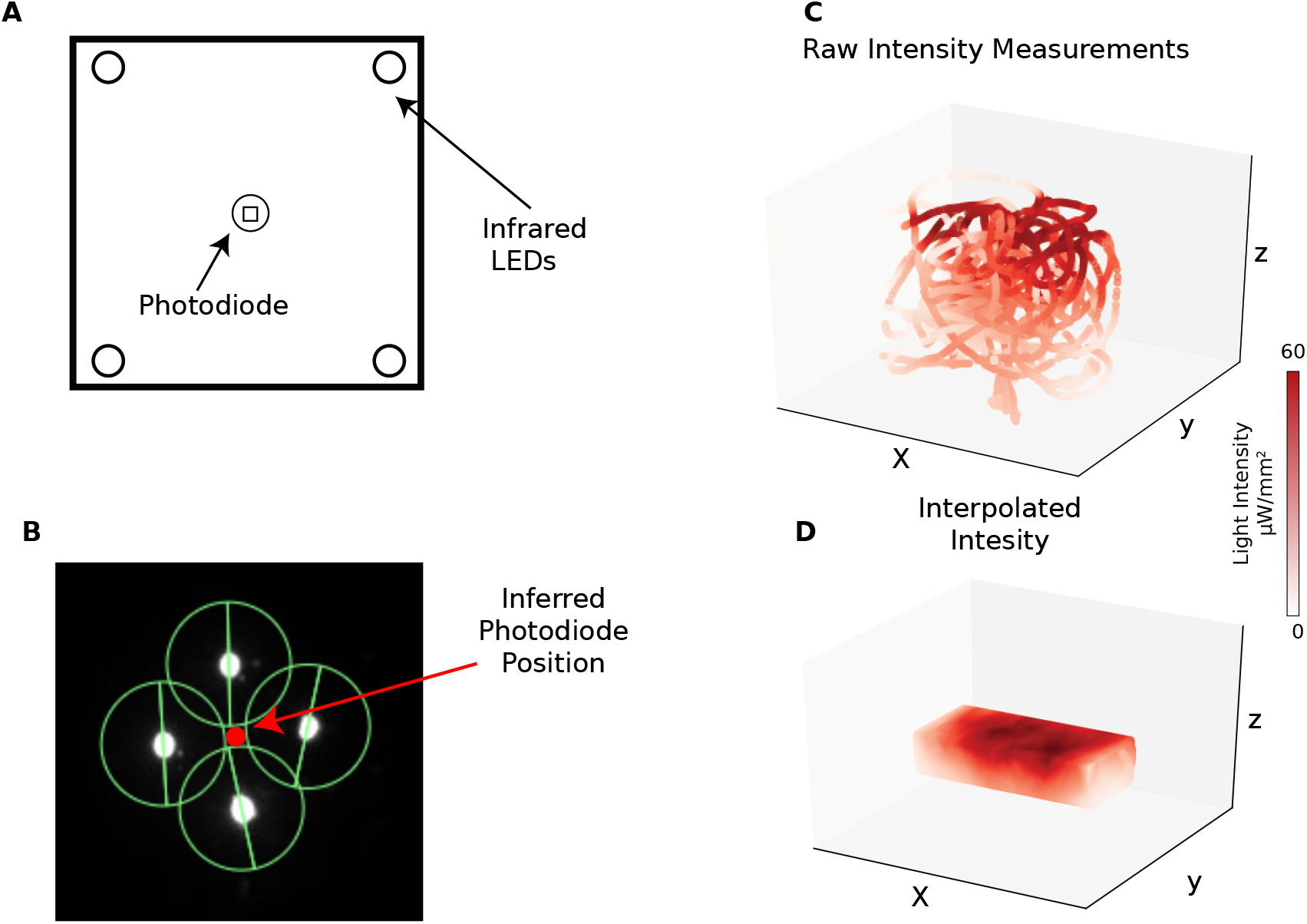
Three dimensional calibration of red light intensity in the wind tunnel volume. Related to Figure 1A. (A) Schematic of light measurement module with four infrared LEDs for tracking and a photodiode in the middle. (B) Example of tracking on four LED’s with inferred position of the photodiode position from the infrared LEDs. (C) Raw position and light intensity measurements from synchronized tracking and photodiode recordings. (D) Interpolated light intensity in the trigger volume.

**Figure S2.**
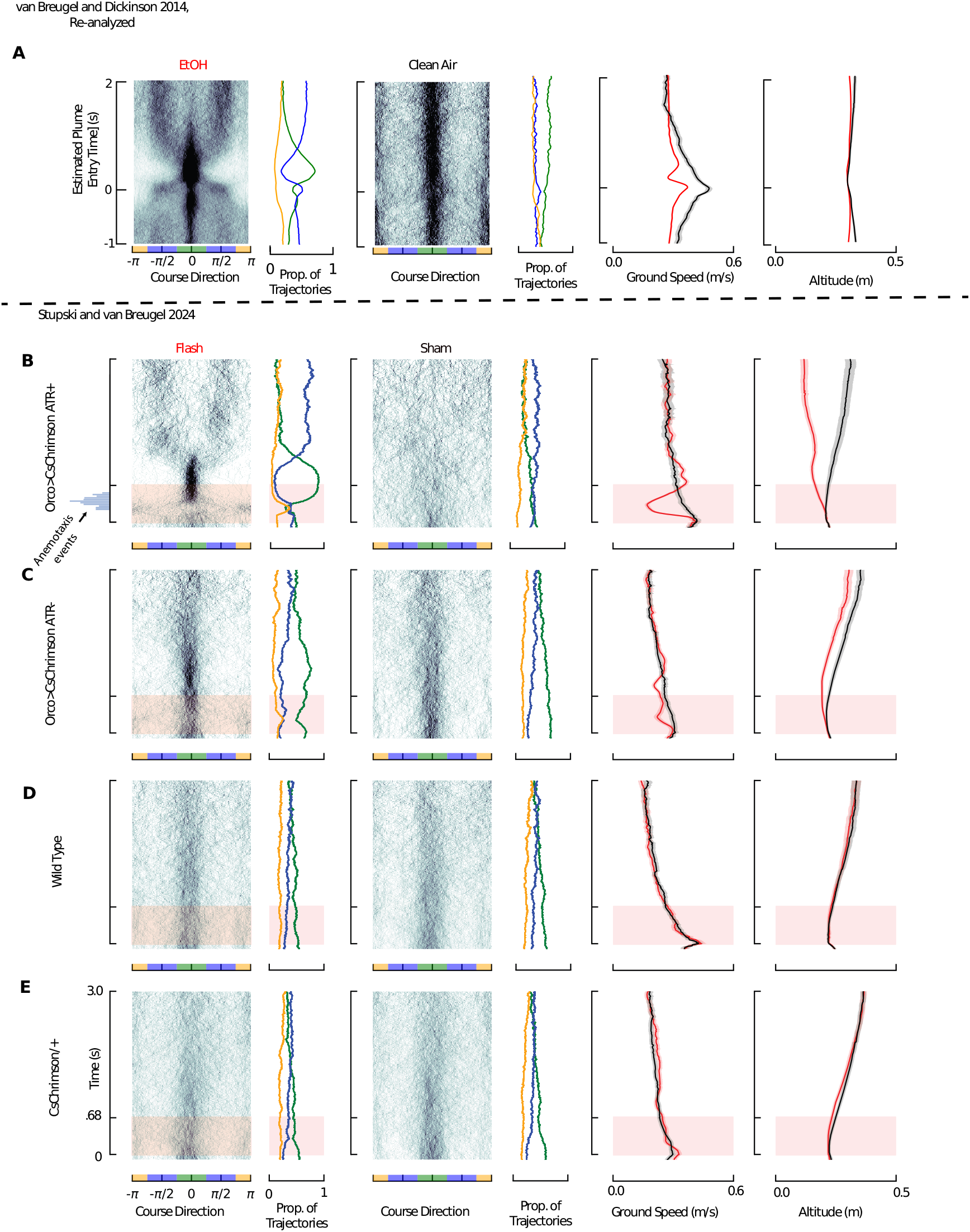
Comparison of Orco>CsChrimson optogenetic activation to wild-type flies encountering an ethanol plume, and genetic controls, all in laminar wind. Related to Figure 1C-F. (A) Reanalysis of data from van Breugel and Dickinson^8^ where wild-type flies encountered a narrow plume of ethanol. Here we analyze the data using the same time scales and graphical parameters as our Orco>CsChrimson experiments shown in Figure 1D and repeated here in panel (B). For the flies encountering ethanol, instead of “flashes” and “shams”, analyses are generated from the separate ethanol (red trace, n=2487) and clean air (black trace, n=942) experiments. (B) Orco>CsChrimson data, repeated from Figure 1d to aid direct visual comparison to other data sets (N=60 flies:n=232 flash trajectories, n=222 sham trajectories). (C) ATR-control. Orco>CsChrimson flies in the same behavioral paradigm as panel B, except these flies were reared without ATR enriched media (N=30 flies: n=357 flash trajectories, n=350 sham trajectories). (D) Wild-type Flies. Repeated from Figure 1d to aid direct visual comparison (N=75 flies: n=398 flash, n= 429 sham). (E) UAS-CsChrimson/+ heterozygotes subject to the same paradigm used in panels B-E. Used to establish that there were no salient visually mediated artifacts from red light flashing associated with *White* heterozygosity (N=75 flies: n=682 flash trajectories, n=683 sham trajectories).

**Figure S3.**
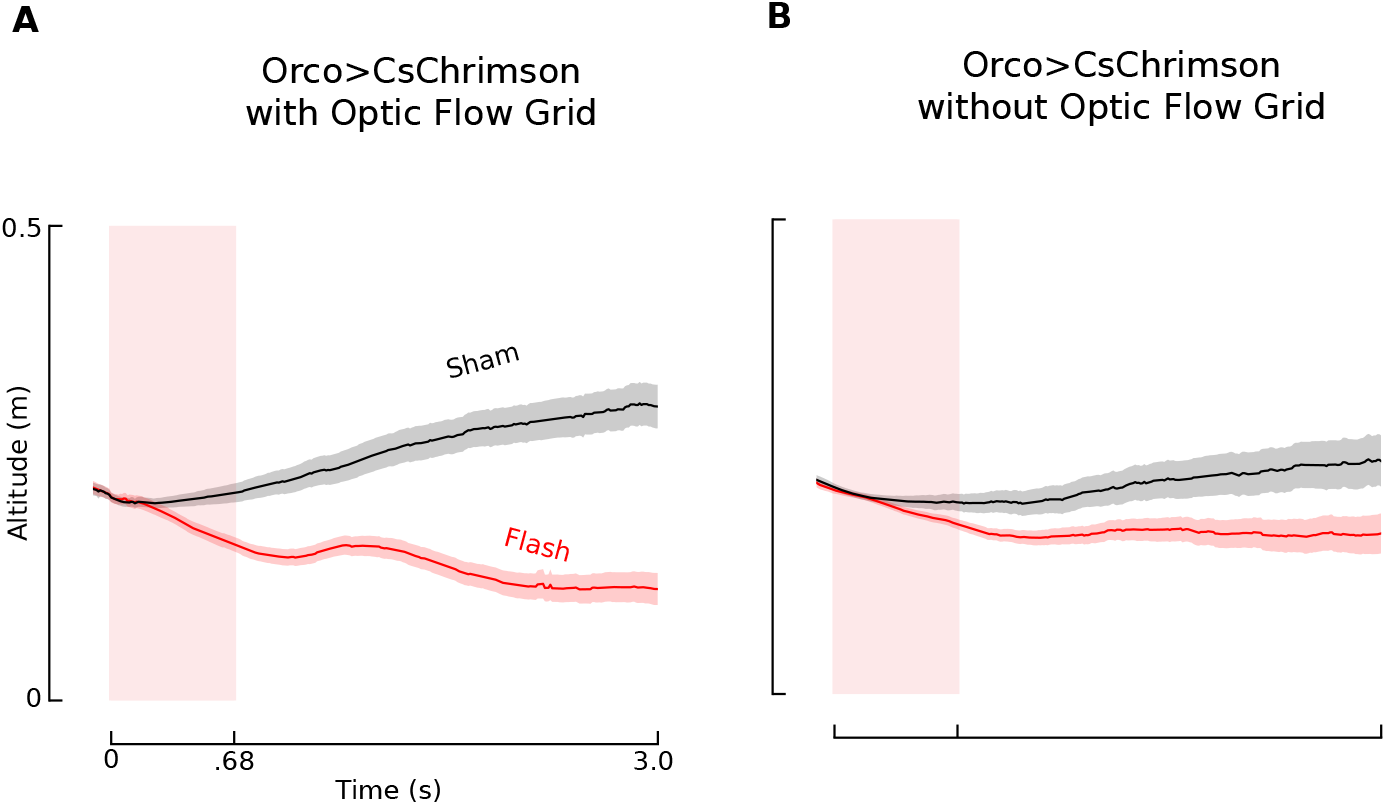
Flies reduce altitude after virtual odor stimulus regardless of the visual contrast on the floor of the wind tunnel in still air. Related to Figure 2E. (A) Altitude dynamics of Orco>CsChrimson flies with a high contrast checkerboard grid on the floor of the wind tunnel (N=60 flies:n= 263 flash trajectories; n=291 sham trajectories). (B) Same as A but without a checkerboard grid on the floor of the tunnel (N=30 flies:n= 740 flash trajectories, n= 786 sham trajectories).

**Figure S4.**
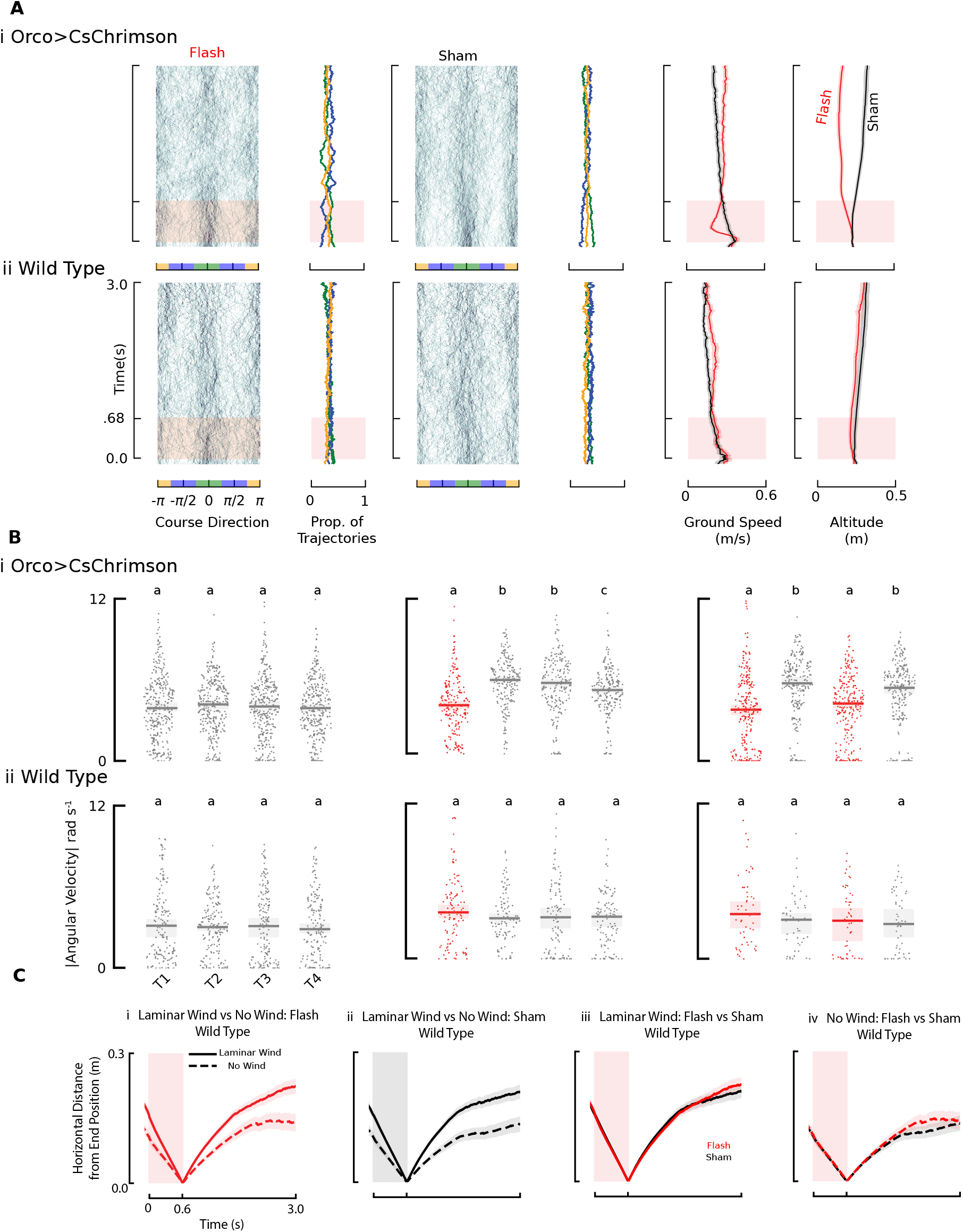
Wild-type controls for still air experiments. Related to Figures 2C-E, 4B-C, 5C. (Ai) Orco>CsChrimson flies repeated from Figure 1c to aid in direct comparisons (N=60 flies:n=263 flash; n=291 sham). (Aii) Wild-type control data for no wind flash and sham conditions (N=135 flies: n=187 flash trajectories; n=184 sham trajectories).(B) Wild-type control for double flash experiments in still air. (Bi) Repeated from Figure 4C to aid in direct comparison of the data. (Bii) Wild-type control for double pulse experiment, analyzed and plotted as in Figure 4C for the sham condition (N=225 flies: n=163 trajectories), single flash condition (N=135 flies:n=107 trajectories), or the double flash condition (N=90 flies: n=50 trajectories). Letters indicate statistical groupings within a genotype and stimulus regimen (Mann-Whitney U test, Bonferroni corrected *α* = .0083). (C)Wild-type control for proximal search analysis. (Ci) Distance from stimulus end position between wild-type fly trajectories which received a light flash in laminar wind : solid line,N=75 flies: n=398 trajectories) or still air (dashed line,N=135 flies: n=187 trajectories). (Cii) same as **a** but for sham trajectories (laminar wind, N=75 flies: n= 429 trajectories; still air, N=135 flies: n= 184 trajectories) (Ciii) Side by side comparison for wild-type trajectories in laminar wind which received either a flash (red) or a sham (black). (Civ) Same except for still air.

**Figure S5.**
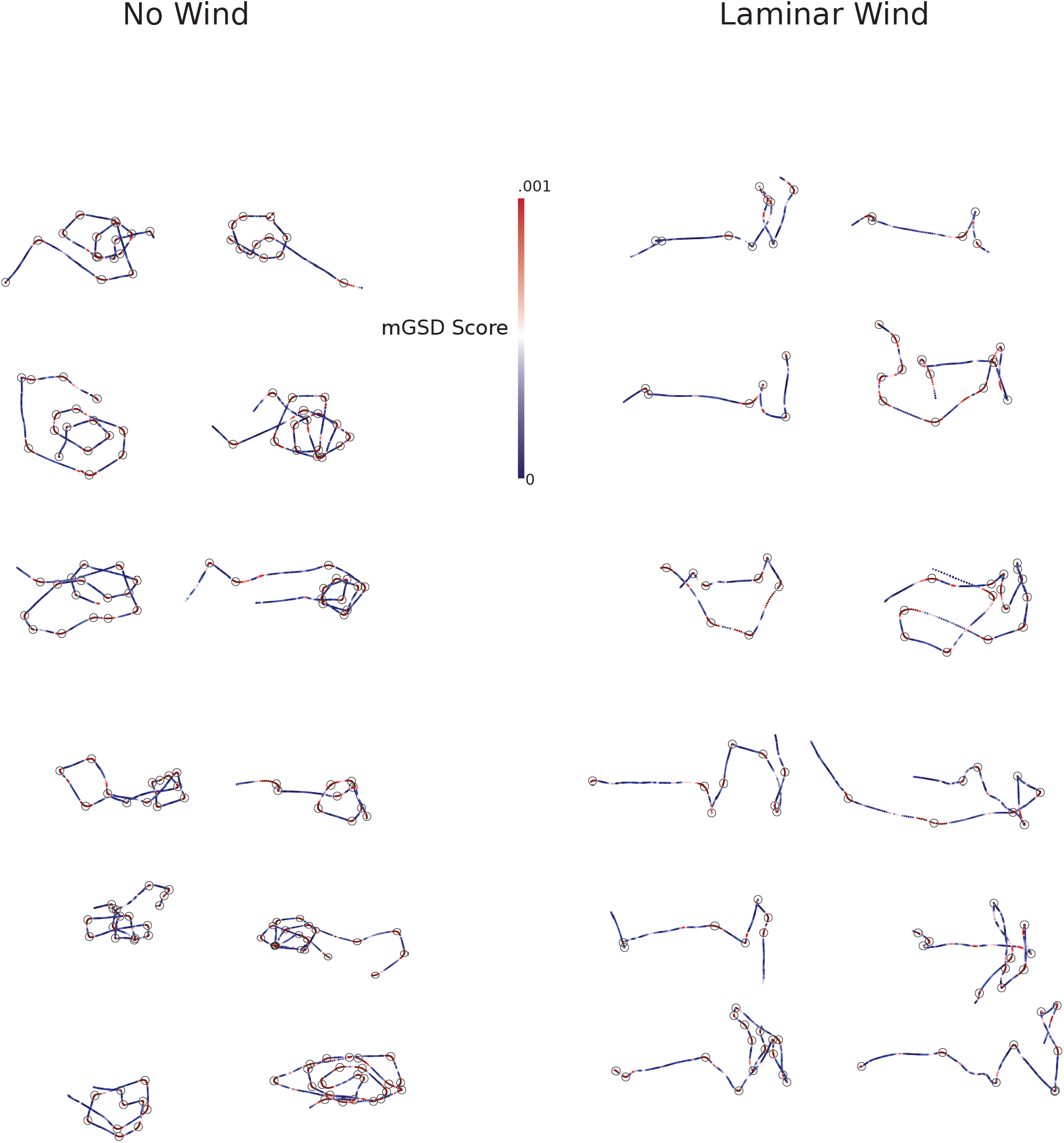
mGSD based saccade labelling. Related to Figure 3A-D. Sample trajectories of Orco>CsChrimson flies which received an optogenetic stimulus (12 from still air conditions (Left) and 12 from laminar wind conditions (Right) color coded by the mGSD score at each point. Black circles indicate locations of saccades as determined by the mGSD algorithm.

**Figure S6.**
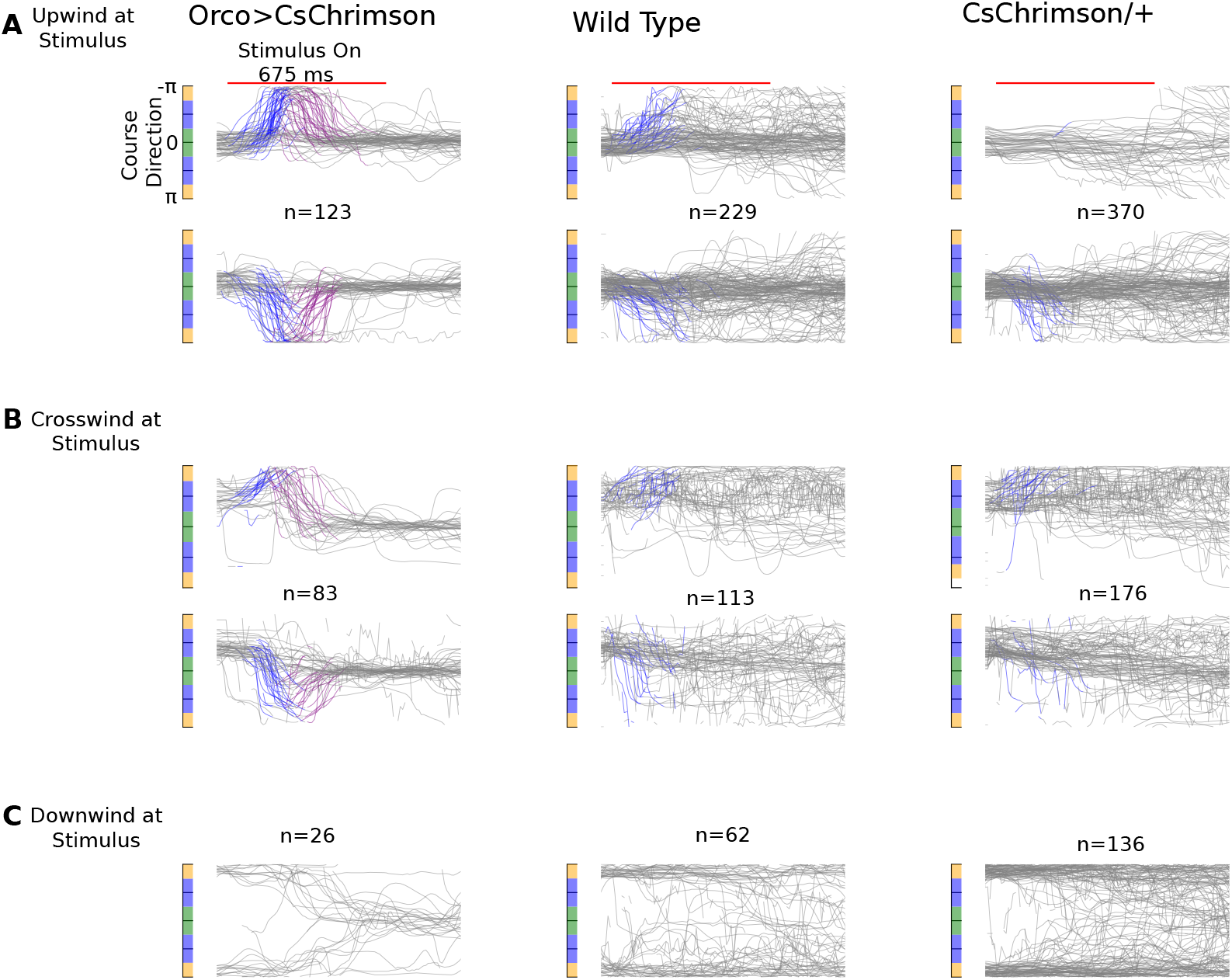
Course changes in laminar wind after light flash are not a visual startle response. Related to Figure 6A-C. Direction vs. time plots for Orco>CsChrimson flies, wild-type flies, and CsChrimson/+ heterozygotes which received a flash event. Flies are separated into those oriented (A) upwind at the time of the stimulus, (B) crosswind at the time of the stimulus, or (C) downwind at the time of the stimulus. For wild-type and CsChrimson/+ flies, blue regions include trajectories that were sufficiently large to qualify as an “anemometric” turn in our analysis. Downwind oriented flies are only shown in gray.

**Figure S7.**
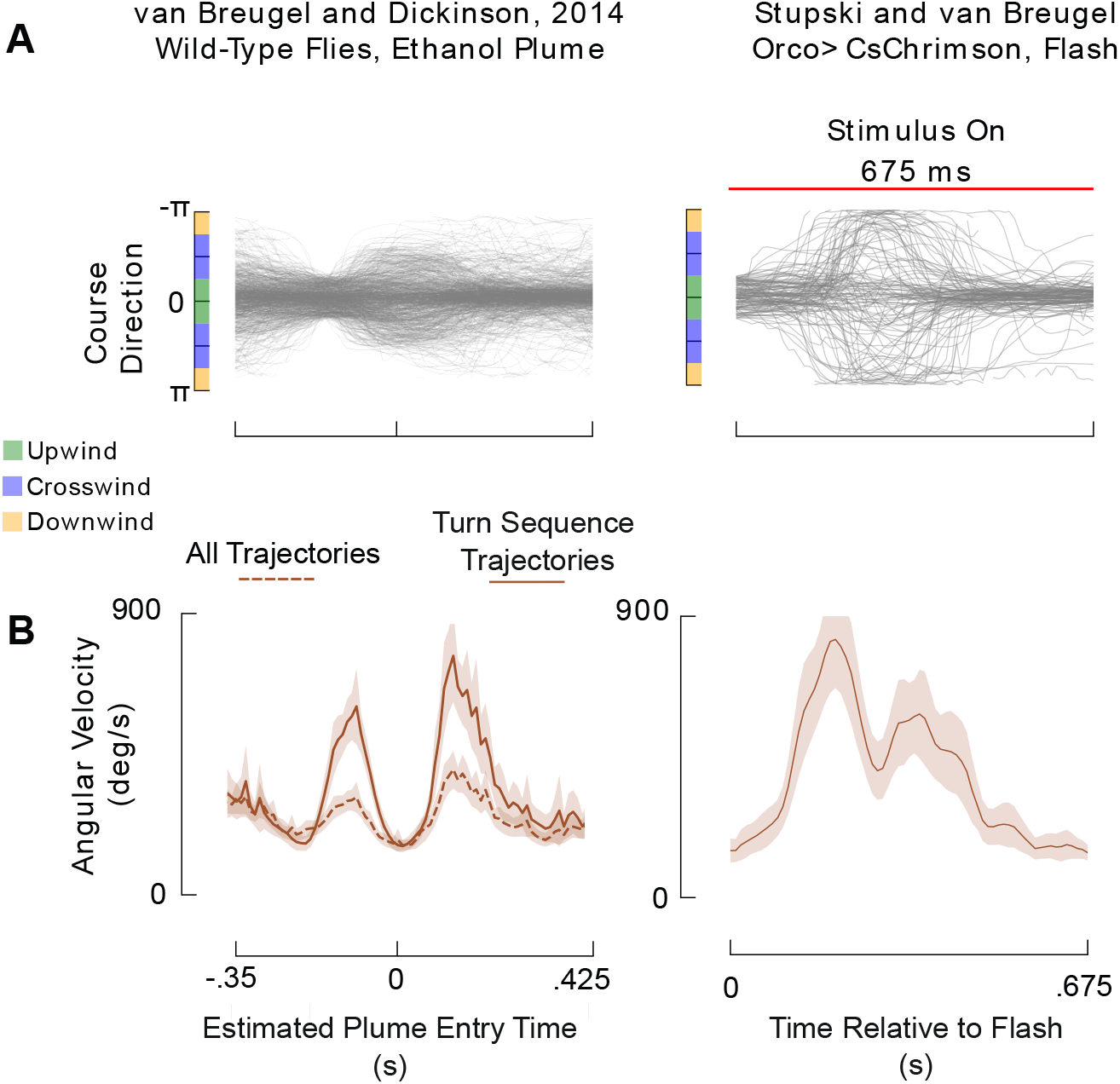
Upwind-oriented flies often perform two-turns after encountering an ethanol plume in laminar wind. Related to Figure 6A-C. (A) Left, course direction vs. estimated time of ethanol plume entry, data from van Breugel and Dickinson^8^. Here we include only trajectories that were oriented upwind 200 ms before the estimated time of plume entry, n=838 plume entry events. Right, Orco>CsChrimson flies which received a flash event when they were oriented upwind, n=123, repeated from Figure 6C. In Figure 6C we identified a two-turn sequence in 85 of these trajectories (69%) (B) Angular velocity vs. time for flies encountering an ethanol plume (left) for both the the whole population (dashed, n=838 plume entry events) and the subset of the population where the two-turn sequence was detected (solid, n=315 plume entry events). Thus, in the ethanol datasets 37% of flies exhibited the two-turn sequence. (Right) Orco>CsChrimson flies receiving a flash. Our reanalysis of the data from van Breugel and Dickinson^8^ suggests that the true estimated plume entry time may have been ∼ 200 ms prior to the originally estimated plume entry time.

## Footnotes

1 Note: flies could determine the airflow magnitude using their antennae. Alternatively, flies could measure the forces on their wings using strain sensors, or use the level of power muscle activity together with an internal model of how that level of activity translates to wing forces.

## Notes

### Competing Interest Statement

The authors have declared no competing interest.

### Summary of Updates

Primary revisions include the addition of a new summary figure, and more detailed theoretical foundation for wind presence estimation.

